# Loss of soluble guanylyl cyclase in platelets contributes to atherosclerotic plaque formation and vascular inflammation

**DOI:** 10.1101/2021.11.15.467680

**Authors:** Carina Mauersberger, Hendrik B. Sager, Jana Wobst, Tan An Dang, Laura Lambrecht, Simon Koplev, Marlène Stroth, Noomen Bettaga, Jens Schlossmann, Frank Wunder, Andreas Friebe, Johan L.M. Björkegren, Lisa Dietz, Peter Sandner, Oliver Soehnlein, Heribert Schunkert, Thorsten Kessler

**Affiliations:** German Heart Centre Munich, Department of Cardiology, Technical University of Munich, Munich, Germany; German Centre for Cardiovascular Research (DZHK e.V.), partner site Munich Heart Alliance, Munich, Germany; Department of Genetics and Genomic Sciences, Icahn Institute for Genomics and Multiscale Biology, Icahn School of Medicine at Mount Sinai, New York, NY, USA; Cancer Research UK Cambridge Institute, University of Cambridge, Li Ka Shing Centre, Cambridge, United Kingdom; Department of Pharmacology and Toxicology, University of Regensburg, Regensburg, Germany; Bayer AG, R&D Pharmaceuticals, Wuppertal, Germany; Institute of Physiology, Julius Maximilian University of Würzburg, Würzburg, Germany; Integrated Cardio Metabolic Centre, Department of Medicine, Karolinska Institutet, Karolinska Universitetssjukhuset, Huddinge, Sweden; Department of Physiology, Institute of Biomedicine and Translational Medicine, University of Tartu, Tartu, Estonia; Institute for Cardiovascular Prevention, Ludwig Maximilian University of Munich, Munich, Germany; Institute for Experimental Pathology, University of Münster, Münster, Germany; Department of Physiology and Pharmacology (FyFa) and Department of Medicine, Karolinska Institutet, Stockholm, Sweden

**Keywords:** Atherosclerosis, Genetics, Soluble guanylyl cyclase, Platelets, Drug treatment

## Abstract

**Aim:** The role of platelets in atherosclerosis remains incompletely understood. Variants in genes encoding the soluble guanylyl cyclase (sGC) in platelets are associated with coronary artery disease (CAD) risk. Here we sought to investigate the contribution of platelet sGC to atherosclerosis and the therapeutic potential of targeting sGC in atherosclerosis.

**Methods and Results:** We genetically deleted sGC in platelets of atherosclerosis-prone *Ldlr*^-/-^ mice. By intravital fluorescence microscopy such Pf4-*Cre*^+^*Gucy1b1*^flox^/^flox^*Ldlr*^-/-^ mice displayed enhanced leukocyte adhesion to atherosclerotic plaques in comparison with their litter mates. Moreover, histological and flow cytometry analyses revealed more numerous inflammatory leukocytes and larger plaque sizes in aortic tissue of *Ldlr*^-/-^ mice lacking sGC in platelets. *In vitro*, supernatant from activated platelets lacking sGC promoted leukocyte adhesion to endothelial cells (EC) via enhanced EC activation. Using cytokine profiling, we identified reduced angiopoietin-1 release by Pf4-*Cre*^+^*Gucy1b1*^flox^/^flox^ and human *GUCY1A1* risk allele carrier platelets to be responsible for enhanced activation of EC and subsequent leukocyte adhesion. Pharmacological sGC stimulation increased platelet angiopoietin-1 release *in vitro* and reduced recruitment of adoptively transferred leukocytes in *Ldlr*^-/-^ mice fed a Western diet. Pharmacological sGC stimulation further reduced atherosclerotic plaque formation and vascular inflammation.

**Conclusion:** Loss of sGC in platelets contributes to atherosclerotic plaque formation via reduced release of the soluble factor angiopoietin-1 and, subsequently, enhanced leukocyte recruitment. Pharmacological sGC stimulation might represent a novel therapeutic strategy to prevent and treat CAD.

**Translational perspective:** Reduced platelet soluble guanylyl cyclase activity contributes to atherosclerotic plaque formation and vascular inflammation. Stimulators of the soluble guanylyl cyclase, an emerging class of drugs already used in pulmonary hypertension and heart failure, are able to reduce atherosclerosis and inflammation in this preclinical model. Together with evidence from human genetics, our findings suggest a promising role of soluble guanylyl cyclase stimulation to prevent coronary artery disease.

## Introduction

Coronary artery disease (CAD) is a leading cause of morbidity and mortality in industrialized nations ^1^. Within the past 15 years, large-scale genetic studies led to the identification of more than 160 genomic variants that are associated with CAD and myocardial infarction (MI) risk. Interestingly, most of the affected genes do not influence traditional risk factors ^2^ but rather mechanisms not previously implicated in the pathophysiology of atherosclerosis. One prominent example is the nitric oxide- (NO-) cyclic guanosine monophosphate- (cGMP-) signaling pathway, which contains several genome-wide significantly associated variants for CAD ^3^. In the center of this pathway sits the soluble guanylyl cyclase (sGC) which is activated by NO and produces the second messenger cGMP. From a genetic perspective private mutations in the genes encoding for sGC as well as rare coding and common non-coding variants in the *GUCY1A1* gene, which encodes the α1-subunit of the sGC, were all associated with CAD or premature MI by exome-sequencing and genome-wide association studies, respectively ^4,5^. These mutations and variants reduce sGC expression or activity ^5-7^; in line, enhanced NO-cGMP-signaling has been associated with reduced risk of several cardiometabolic phenotypes including CAD and peripheral artery disease ^8^.

*GUCY1A1* and *GUCY1B1*, genes encoding sGC, are expressed at high levels in platelets and vascular smooth muscle cells. In humans, it was shown that carriers of the CAD risk variant rs7692387 have reduced sGC-α_1_ protein levels, which might impair the effects of the natural platelet inhibitor NO ^7^. Indeed, a retrospective analysis of two randomized trials revealed that inhibition of platelet activity by aspirin successfully reduced cardiovascular events in primary prevention only in homozygous carriers of the *GUCY1A1* risk allele. Since the overall role of platelets in atherosclerosis remains controversial ^9,10^, we decided to delete sGC in mice platelet-specifically by knockout of its β_1_-subunit in order to investigate the contribution of platelet sGC on atherosclerosis and vascular inflammation and to further evaluate the potential of sGC stimulation as a therapeutic strategy.

## Methods

### Mouse studies

Animal experiments were conducted in accordance with the German legislation on protection of animals and approved by the local animal care committee (Az: ROB-55.2-2532.Vet_02-15-176). Platelet-specific sGC knockout mice (Pf4-*Cre*^*+*^*Gucy1b1*^flox/flox^) were obtained by crossing conditional NO-GC β1 knockout mice (*Gucy1b1*^flox/flox^) ^11^ and Pf4-*Cre*^*tg/+*^ mice (Pf4-*Cre*^*+*^) as previously described ^12^. Subsequently, Pf4-*Cre*^*+*^*Gucy1b1*^*flox/flox*^ mice were crossbred with *Ldlr*^*-/-*^ mice (B6.129S7-*Ldlr*^tm1Her^/J) to induce a *Ldlr*^*-/-*^ pro-atherosclerotic background. For experiments, Pf4-*Cre*^*+*^*Gucy1b1*^*flox/flox*^*Ldlr*^-/-^ and *Gucy1b1*^*+/flox*^*Ldlr*^-/-^ mice were mated to receive control (Pf4-*Cre*^*+*^*Gucy1b1*^+^/^flox^*Ldlr*^-/-^) and platelet-specific sGC knockout mice (Pf4-*Cre*^*+*^*Gucy1b1*^*flox/flox*^*Ldlr*^-/-^) littermates. *Irag1*^flox/flox^ mice ^13^ were mated with Pf4-*Cre*^*+*^ mice and obtained Pf4-Cre^+^*Irag1*^flox/flox^ mice were backcrossed to a C57BL/6J background. Pf4-*Cre*^+^*Irag1*^+/flox^ and Pf4-*Cre*^+^*Irag1*^+/+^ littermates were used as controls. *Ldlr*^*-/-*^ and *Ubc-GFP* (C57BL/6-Tg(UBC-GFP)30Scha/J) mice were purchased from The Jackson Laboratory (Bar Harbor, ME, USA). Only male or male and female mice at a 1:1 ratio at 8 to 12 weeks of age were used for all experiments.

To compare aortic plaque sizes between Pf4-*Cre*^*+*^*Gucy1b1*^*flox/flox*^*Ldlr*^-/-^ and Pf4-*Cre*^*+*^*Gucy1b1*^+/flox^*Ldlr*^-/-^ mice (in this comparison termed *Ldlr*^-/-^), animals were fed a western diet (21.2 % fat and 0.2 % cholesterol by weight; TD.88137; Envigo, Indianapolis, IN) for ten weeks. For intravital fluorescence microscopy, Pf4-*Cre*^*+*^*Gucy1b1*^*flox/flox*^*Ldlr*^-/-^ and Pf4-*Cre*^*+*^*Gucy1b1*^+/flox^*Ldlr*^-/-^ mice (in this comparison termed *Ldlr*^-/-^) were fed a western diet (21.2 % fat and 0.2 % cholesterol by weight; TD.88137; Envigo, Indianapolis, IN) for six weeks. To compare aortic plaque sizes after pharmacological sGC stimulation, *Ldlr*^*-/-*^ mice were fed a western diet (21.1 % crude fat and 0.15 % cholesterol, E15721-34; TD.88137 modified; ssniff, Soest, Germany) containing either 0 (control group) or 150 ppm (treatment group) of the sGC stimulator BAY-747 (N-(2-amino-2-methylbutyl)-8-[(2,6-difluorobenzyl)oxy]-2,6-dimethylimidazo[1,2-a]pyridine-3-carboxamide) (Bayer AG, Leverkusen, Germany) ad libitum for ten weeks. For adoptive transfer experiments, *Ldlr*^*-/-*^ mice were fed the same diet for six weeks.

### Human samples

The study protocol was approved by the local ethics committee of the Technical University of Munich (387/17 S). Blood was collected from healthy volunteers after signing the informed consent. To determine the genotype of the subjects, DNA was isolated from whole blood using the Gentra Puregene Blood Kit (Qiagen, Hilden, Germany) according to the manufacturer’s protocol. Samples were genotyped for the *GUCY1A1* risk variant using a rs7692387 TaqMan Genotyping Assay (C_29125113_10) on a Viia7 system (both Thermo Fisher Scientific, Waltham, MA, USA).

### Histology, immunohistochemistry and en face staining

Aortic roots were embedded in optimal cutting temperature compound (Sakura Finetek, Tokyo, Japan) and snap frozen to -80 °C. Frozen samples were cut into 5 μm sections and applied to microscope slides. From the onset of aortic valves, every fifth slide was subjected to tissue staining. For Masson Trichrome stain, the procedure of Masson as modified by Lillie was applied according to manufacturer’s instructions (Sigma Aldrich, St. Louis, MO, USA). In brief, frozen sections were hydrated and fixated in 4 % paraformaldehyde before mordanting in Bouin’s solution (Sigma Aldrich) at 56 °C for 15 minutes. Afterwards, cell nuclei were stained in Weigert’s Iron Hematoxylin Solution (Sigma Aldrich) and darkened in running tap water. Specimens were successively subjected to Biebrich Scarlet-Acid Fuchsin solution, phosphotungstic/ phosphomolybdic acid solution and aniline blue solution for staining of cytoplasm, muscle and collagen structures, respectively. After rinsing in 1 % acetic acid, slides were dehydrated in an increasing ethanol row followed by xylene and finally mounted in Mountex (MEDITE, Burgdorf, Germany). Mean total plaque size (in μm^2^) was evaluated for sections showing at least two complete cusps and analyzed using ImageJ.

For immunohistochemistry, specimens were fixated in ice-cold acetone, blocked in 10 % rabbit serum in phosphate-buffered saline (PBS) with Tween 20 (0.2 %; all Sigma Aldrich) and stained in anti-CD11b antibody (101202, BioLegend, San Diego, CA, USA) or anti-monocyte + macrophage (MoMa) antibody (ab33451, abcam, Cambridge, MA, USA) overnight, followed by incubations in HRP conjugated secondary antibody (ab6734) and AEC substrate (ab64252, both abcam). Subsequently, cell nuclei were counterstained in Gill’s hematoxylin solution II (Merck Millipore, Darmstadt, Germany). CD11b or monocyte and macrophage content was quantified as positive area per total plaque area using ImageJ software.

For *en face* analyses, aortae were dissected from the heart to the iliac bifurcation, cleaned of surrounding tissue and fixed for 24 hours at 4°C in a 4% solution of paraformaldehyde in PBS. Samples were washed in 60% isopropanol and incubated for 30 minutes at 37°C in a freshly filtered solution of 3 mg/ml oil red O (O0625, Sigma Aldrich) in 60% isopropanol. After washing off excess dye in 60% isopropanol, aortae were opened longitudinally, pinned on a black pad and imaged on a Stemi 2000-C microscope with an Axiocam ERc 5s camera using ZEN 2 software (Carl Zeiss, Oberkochen, Germany). Total area and lesion area of the whole aorta from aortic root to branch of the right renal artery were determined using Image J.

### Generation of supernatant from activated platelets

800 μl each of blood from Pf4-*Cre*^+^*Gucy1b1*^+/flox^ mice and Pf4-*Cre*^*+*^*Gucy1b1*^flox/flox^ mice was collected in heparinized tubes, gently diluted in PBS and centrifuged for 10 minutes at 100 g and room temperature without active deceleration. Platelet rich plasma was subjected to a second centrifugation step at 700 g to obtain platelets which were resuspended in RPMI 1640 (Thermo Fisher Scientific) and activated by orbital shaking for 30 minutes at 1000 rpm. Samples were centrifuged at 12,000 g for 10 minutes and the supernatant was directly used in subsequent experiments. Thrombocyte count was analyzed simultaneously with an automated hematology analyzer (Sysmex Corporation, Kobe, Japan).

For sGC stimulation, platelet suspension of wildtype (WT) mice was split in two vials containing either BAY-747 (150 ppm (150 mg/l) final concentration) or vehicle (DMSO, 0.2 %), mixed gently and incubated for 30 minutes prior to shaking.

Blood from healthy volunteers was collected in hirudin-coated tubes (Sarstedt, Nümbrecht, Germany) and centrifuged for 13 minutes at 170 g and room temperature without active deceleration. Platelet rich plasma was transferred into new tubes and activated by orbital shaking for 30 minutes at 1000 rpm. Afterwards, samples were centrifuged at 12,000 g for 10 min and the supernatant was stored at -80°C before conducting subsequent experiments. Thrombocyte count was determined from platelet rich plasma as stated above.

### In vitro leukocyte adhesion

Bone marrow monocytes and neutrophils were isolated from C57BL/6J mice using MACS cell separation columns (Miltenyi Biotec, Bergisch Gladbach, Germany) after incubation with either anti-Ly6G-PE (clone 1A8) or anti-CD115-Biotin (clone AFS98, both BioLegend) antibodies followed by PE- and streptavidin-coated microbeads (Miltenyi), respectively.

Primary murine aortic endothelial cells (C57-6052, mAoEC, Cell Biologics, Chicago, IL, USA) were cultured in complete endothelial cell medium (PB-M1168, PeloBiotech, Planegg, Germany) in a humidified incubator with 5 % CO_2_ at 37°C and grown to confluency in 96-well-plates for experiments.

Adhesion assays were performed using the CytoSelect Leukocyte-Endothelium Adhesion Assay (Cell Biolabs, San Diego, CA, USA) according to the manufacturer’s instructions. Briefly, leukocytes were fluorescently labelled with LeukoTracker solution, resuspended in RPMI 1640 (Thermo Fisher Scientific) and added to mAoECs to a number of 2.5 × 10^5^ cells. Cells were incubated for one hour at 37°C in the presence of 50 μl of activated platelet supernatant. Plates were washed trice to remove non-adherent cells, lysed in 1X lysis buffer and fluorescence was measured on an Infinite M200 PRO plate reader (TECAN, Männedorf, Switzerland; Exc = 485 nm and Em = 535 nm). Experiments were conducted in triplicates.

For preincubation experiments, either neutrophils or endothelial cells were exclusively incubated with activated platelet supernatant of WT mice for 30 minutes prior to performing the adhesion assay, omitting the additional administration of platelet supernatant in this step. To inhibit Tie2-receptor effects, mAoECs were preincubated with 0.5 μM BAY-826 (R&D Systems, Minneapolis, MN, USA) or vehicle (DMSO, 0.1 %) for five hours prior to performing the adhesion assay.

### Intravital fluorescence microscopy and in vivo leukocyte adhesion

Pf4-*Cre*^*+*^*Gucy1b1*^+/flox^*Ldlr*^-/-^ (in this comparison termed *Ldlr*^-/-^) mice and Pf4-*Cre*^*+*^*Gucy1b1*^flox/flox^*Ldlr*^-/-^ mice were anaesthetized with a combination of midazolam, medetomidine and fentanyl and subjected to intravital microscopy of the right carotid artery bifurcation as described previously ^14^. Leukocytes were labelled *in vivo* by intravenous injection of anti-Ly6G (PE conjugated, clone 1A8, BioLegend, San Diego, CA, USA), anti-Ly6C (Alexa Fluor 488 conjugated, clone HK1.4), and anti-CD11b (PE-conjugated, clone M1/70, BioLegend) antibodies. To examine leukocyte to endothelium interactions, movies of 30 seconds each were acquired using an Olympus BX51 microscope (Tokyo, Japan) with a Hamamatsu 9100-02 EMCCD camera (Hamamatsu, Japan) and a 10x saline-immersion objective. In subsequent analysis of the movies, number of adherent neutrophils (Ly6G-PE^+^), monocytes (Ly6C-Alexa Fluor 488^+^), and myeloid cells (CD11b-PE^+^) was examined in a blinded manner, considering cells as adherent if their position did not change during imaging.

### Isolation of nucleic acids and (quantitative) polymerase chain reaction

mAoEC were grown to confluence in 12-well dishes and stimulated with activated platelet supernatant from either Pf4-*Cre*^*+*^sGC^+/flox^ (in this comparison termed WT) or Pf4-*Cre*^*+*^sGC^flox/flox^ mice in RPMI 1640 for one hour at 37°C. Cells were lysed by adding 500 μl of TRIzol (Thermo Fisher Scientific) and stored at -80°C.

After adding chloroform (288306, Sigma Aldrich), samples were shaken vigorously and centrifuged at 12,000 g for 15 minutes and 4°C. The upper phase containing RNA was further processed using the RNeasy Mini Kit (Qiagen, Hilden, Germany) according to the manufacturer’s recommendations. RNA was quantified using a NanoQuant Plate on an Infinite M200 PRO plate reader (TECAN) and RNA integrity was measured on a 2100 Bioanalyzer (Agilent Technologies, Santa Clara, CA).

After the RNA was transcribed into cDNA using the High-Capacity RNA-to-cDNA kit (Applied Biosystems, Waltham, MA), real-time quantitative PCR was performed using TaqMan Fast Universal PCR Master Mix (4366072) and TaqMan probes (*Vcam1*, Mm01320970_m1; *Gapdh*, Mm99999915_g1; both Thermo Fisher Scientific). Reactions were performed in a total volume of 10 μl on a ViiA 7 system (Thermo Fisher Scientific). *Gapdh* was used as a housekeeping gene and data were evaluated by conversion to ΔCt values.

### Cytokine profiling and enzyme-linked immunosorbent assays

Proteome profiling assays were performed using the Proteome Profiler Mouse XL Cytokine Array (ARY028, R&D Systems) according to the manufacturer’s protocol. Briefly, after isolating and activating platelets of Pf4-*Cre*^*+*^*Gucy1b1*^+/flox^ mice and Pf4-*Cre*^*+*^*Gucy1b1*^flox/flox^ mice by shaking in RPMI as described above, samples were added to the supplied antibody-spotted nitrocellulose membrane and incubated at 4°C overnight. Captured proteins were detected using a mixture of biotinylated detection antibodies followed by streptavidin-horseradish peroxidase and visualized using chemiluminescent detection reagents. Signal intensities were detected by an ImageQuant LAS 4000 imaging system (GE Healthcare Life Sciences, Freiburg, Germany) and analyzed using the appropriate image analysis software (ImageQuant TL; GE Healthcare Life Sciences). Signal intensities of target proteins were normalized to signal intensities for reference spots in each sample. The procedure was repeated for a total of 6 samples per group.

Angiopoietin-1 enzyme-linked immunosorbent assays (ELISA) were performed to determine murine (EK1296, Boster Biological Technology, Pleasanton, CA) and human (DANG10, R&D systems) protein levels according to the manufacturer’s recommendations.

### Co-expression analyses in STARNET

Aligned multitissue RNA-seq samples from STARNET ^15^ were pseudo log transformed and normalized using L2 penalized regression with penalty term 1.0, adjusting for the covariates: sequencing laboratory, read length, RNA extraction protocol (PolyA and Ribo-Zero), age, and gender. Additional adjustments included the first four surrogate variables detected by Surrogate Variable Analysis (SVA) ^16^ and flow cell information after singular value decomposition retaining components with eigenvalues > 4. Co-expression modules were inferred using weighted gene co-expression network analysis (WGCNA) ^17^ with β=5.2 for tissue-specific and β=2.7 for cross-tissue correlations, resulting in both tissue-specific and cross-tissue co-expression networks as previously described ^18^. These data and analyses were accessed through the STARNET browser (under revision) by querying co-expression modules containing ANGPT1. KEGG pathway and Gene Ontology (GO) enrichment was carried out on co-expression module 11 for transcripts derived from whole blood (1012 out of 1016 transcripts) using Enrichr ^19^.

### Flow cytometry

Mice were sacrificed under isoflurane anaesthesia and blood was collected in EDTA coated microvettes (Sarstedt). For *in vivo* staining of circulating blood leukocytes, an antibody directed against CD45-BV605 (clone 30-F11, 1:10 in 100 μl PBS, BioLegend) was injected intravenously 5 min before euthanizing the animals. After lysing red blood cells in 1X RBC lysis buffer (BioLegend), samples were washed and resuspended in FACS buffer (PBS containing 0.5 % bovine serum albumin, Sigma Aldrich). Aortas were perfused through the left ventricle with PBS and excised from root to common iliac artery bifurcation after removing perivascular fat and surrounding other tissue, minced using fine scissors and digested in 450 U/ml collagenase I, 125 U/ml collagenase XI, 60 U/ml DNase I and 60 U/ml hyaluronidase (Sigma Aldrich) for 1 hour at 37°C under agitation. Cell suspensions were filtered through 40 μm nylon cell strainers (BD Biosciences, San Jose, CA), washed and resuspended in FACS buffer.

Both blood and aortic cells were first stained for hematopoietic lineage markers with phycoerythrin (PE) conjugated anti-mouse antibodies directed against B220 (RA3-6B2, 1:600), CD49b (clone DX5, 1:1200), CD90.2 (clone 53-2.1, 1:3000), NK1.1 (clone PK136, 1:600), Ter119 (clone TER-119, 1:600) and Ly6G (clone 1A8, 1:600) for 15 minutes at 4°C and washed. This was followed by a second round of staining for Ly6C-BV421 (clone HK4.1), CD115-BV510 (clone AFS98), CD45.2-PerCP/Cy5.5 (clone 104, 1:300), F4/80-PE-Cy7 (clone BM8) and CD11b-APC-Cy7 (clone M1/70) in 1:600 dilution, if not stated otherwise. Antibodies were either purchased from BioLegend or BD Biosciences (San Jose, CA).

Cells were submitted to flow cytometric analysis on BD LSRFortessa (BD Biosciences) and analysed with FlowJo software (version 9.9.6, Tree Star, Ashland, OR). Cells were gated on viable (FSC-A vs. SSC-A) and single (FSC-A vs. FSC-W and SSC-A vs. SSC-W) cells. Neutrophils were identified as lineage^high^(CD45.2/CD11b)^high^CD115^low^. Monocytes were identified as lineage^low^(CD45.2/CD11b/CD115)^high^Ly6C^high/low^. Macrophages were identified as lineage^low^(CD45.2/CD11b/F4/80)^high^Ly6C^low/int^.

### Adoptive transfer of leukocytes

Bone marrow monocytes and neutrophils were isolated simultaneously from *Ubc-GFP* mice using anti-Ly6G-PE and anti-CD115-Biotin antibodies followed by PE- and Streptavidin-coated microbeads as described before and resuspended in PBS. We intravenously injected equal amounts of isolated cells into Pf4-*Cre*^*+*^*Gucy1b1*^+/flox^*Ldlr*^-/-^ mice and Pf4-*Cre*^*+*^*Gucy1b1*^flox/flox^*Ldlr*^-/-^ fed a Western diet for 6 weeks and harvested blood and aortae 24 hours later as stated above. The number of CD45.2^high^/CD11b^high^/GFP^high^ cells within the aorta normalized to the exact number of injected cells was quantified by flow cytometry.

### Statistical analysis

Normality distribution of data was assessed using the Kolmogorov-Smirnov test. Test results and subsequently used statistical tests are displayed in **Suppl. Table S1**. Data was analyzed using two-tailed Student’s unpaired or paired t-test (for normally distributed data) or Mann-Whitney test (for non-normally distributed data), as appropriate and indicated in the respective figure legend (and **Suppl. Table S1**). When comparing more than two groups, (RM) one-way ANOVA test followed by a Tukey test for multiple comparisons was performed when data was normally distributed. To determine statistical outliers, the two-sided ROUT’s test was used. If outliers were removed from the analysis, it is indicated in the respective figure legend. Sample sizes/numbers of replicates are indicated in the figure legends and visualized in the figures (each symbol represents one animal/biological replicate) and data are displayed as mean + s.e.m. P-values below 0.05, in case of investigating more than two groups after adjustment for multiple testing, were regarded as statistical significant. Statistical analyses were performed using GraphPad Prism version 9 for macOS (GraphPad Software, La Jolla, CA).

## Results

### Atherosclerotic plaque formation and vascular inflammation in mice lacking platelet sGC

We created mice with a platelet-predominant sGC knockout (Pf4-Cre^+^*Gucy1b1*^flox^/^flox^), which displayed reduced sGC-β_1_ protein levels compared to platelet sGC wild type (WT) mice and – expectedly – reduced inhibition of adenosine diphosphate-induced platelet aggregation by the NO-donor sodium nitroprusside (**Suppl. Fig. S1**). To examine the effect of platelet sGC on atherosclerosis, we crossbred these animals with atherosclerosis-prone mice (*Ldlr*^-/-^ mice) which were then fed a Western diet for ten weeks. Body weight, serum cholesterol and hematologic parameters were comparable between the genotypes (**Suppl. Fig. S2**). Pf4-Cre^+^*Gucy1b1*^flox/flox^*Ldlr*^-/-^ mice displayed larger plaques in the aortic root (246,998 ± 22,162 (n=12) vs. 189,843 ± 15,156 (n=14) μm^2^, p=0.04; **Fig. 1A**) and aorta *en face* analysis (5.4 ± 0.7 (n=8) vs. 3.9 ± 0.2 (n=9) %, p=0.03; **Fig. 1B**). Immunohistochemical stainings of aortic root sections revealed increased plaque monocyte and macrophage content in Pf4-Cre^+^*Gucy1b1*^flox/flox^*Ldlr*^-/-^ compared to *Ldlr*^-/-^ mice (49.1 ± 4.2 (n=12) vs. 38.8 ± 2.8 (n=14) % of plaque area, p=0.04; **Fig. 1C**). To explore why monocyte/macrophages accumulated more in mice with platelet-predominant sGC depletion, we performed intravital fluorescence microscopy of carotid artery plaques in Pf4-Cre^+^*Gucy1b1*^flox/flox^*Ldlr*^-/-^ and *Ldlr*^-/-^ mice which were fed a Western diet for six weeks to initiate plaque formation. We found enhanced adhesion of leukocytes in Pf4-Cre^+^*Gucy1b1*^flox/flox^*Ldlr*^-/-^ compared to *Ldlr*^-/-^ mice (20.9 ± 1.5 (n=13) vs. 13.6 ± 1.4 (n=11) Cd11b^+^ cells, p<0.01; **Fig. 1D, Suppl. Fig. S3**). In line, flow cytometry of aortic cell suspensions revealed more numerous Ly6C^high^ monocytes (1,829 ± 262 vs. 964 ± 129 cells/aorta, n=11 each, p=0.01), neutrophils (974 ± 170 vs. 392 ± 60 cells/aorta, n=11 each, p<0.01), and macrophages (14,895 ± 1,912 vs. 9,463 ± 1,211 cells/aorta, n=11 each, p=0.02; **Fig. 1E**) in Pf4-Cre^+^*Gucy1b1*^flox/flox^*Ldlr*^-/-^ compared to *Ldlr*^-/-^ mice while blood leukocyte numbers remained unchanged (**Suppl. Fig. S4**). Taken together, these data indicate that the lack of sGC in platelets contributes to vascular inflammation and hence atherosclerotic plaque progression.

**Figure 1:**
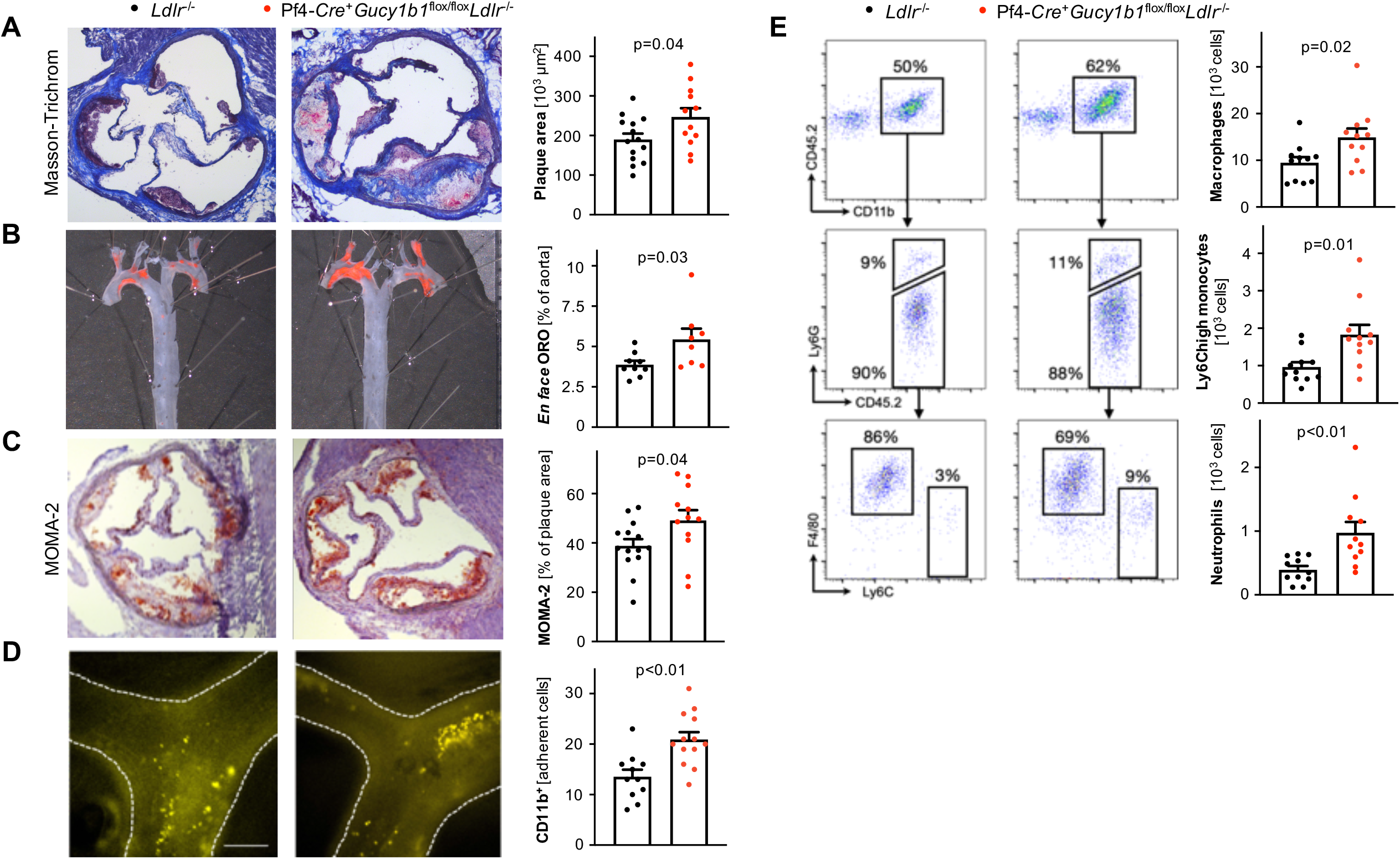
Atherosclerotic plaque formation and vascular inflammation in mice lacking platelet sGC. **A**-**C**. Atherosclerotic plaque formation (**A, B**) and monocyte and macrophage content (**C**) in Pf4-*Cre*^+^*Gucy1b1*^flox/flox^*Ldlr*^-/-^ compared to *Ldlr*^-/-^ mice which were fed a western diet for ten weeks. **D**. Leukocyte adhesion to atherosclerotic plaques in Pf4-*Cre*^+^*Gucy1b1*^flox/flox^*Ldlr*^-/-^ compared to *Ldlr*^-/-^ mice which were fed a western diet for six weeks to induce atherosclerotic plaque formation. Unpaired t-test. E. Quantification of vascular inflammation by flow cytometry analysis of aortic cell suspensions of Pf4-*Cre*^+^*Gucy1b1*^flox/flox^*Ldlr*^-/-^ compared to *Ldlr*^-/-^ mice. Mann-Whitney test. Each symbol represents one animal. Data are mean and s.e.m.

### Platelet sGC and leukocyte adhesion in vitro

To follow-up on enhanced adhesion of leukocytes to atherosclerotic plaques in mice lacking platelet sGC under proathergenic conditions, we tested whether this phenotype can be resembled *in vitro*. Therefore, we isolated blood monocytes and neutrophils from wild type (WT) mice and incubated these with WT aortic endothelial cells (EC) in the presence of supernatant from activated platelets from WT or Pf4-*Cre*^+^*Gucy1b1*^flox/flox^ mice. We found that incubation with the supernatant of activated platelets lacking sGC enhanced leukocyte, particularly monocyte (33,125 ± 1,313 vs. 27,039 ± 555 relative fluorescence units (rfu), p<0.01, n=8 experiments; **Fig. 2A**) and neutrophil (33,810 ± 1,139 vs. 27,824 ± 758 rfu, p<0.001, n=8 experiments; **Fig. 2B**) adhesion. To delineate whether leukocytes or EC are activated by the sGC knockout platelet releasate, we preincubated neutrophils and EC with activated platelet supernatant from Pf4-*Cre*^+^*Gucy1b1*^flox^/^flox^ mice prior to performing the adhesion assay. We found that preincubation of EC with supernatant from activated sGC knockout platelets increased adhesion compared to preincubation of leukocytes (31,383 ± 1,731 vs. 22,254 ± 1,662 rfu, p_adj_<0.001, n=12 experiments; **Fig. 2C**). In line, already at this very early timepoint, we found enhanced expression of the adhesion molecule *Vcam1* in EC that were incubated with supernatant of activated Pf4-*Cre*^+^*Gucy1b1*^flox/flox^ platelets (**Suppl. Fig. S5**). These data indicate i) that platelets from WT and Pf4-*Cre*^+^*Gucy1b1*^flox/flox^ mice differentially release a soluble factor and ii) that this preferentially leads to activation of EC.

**Figure 2:**
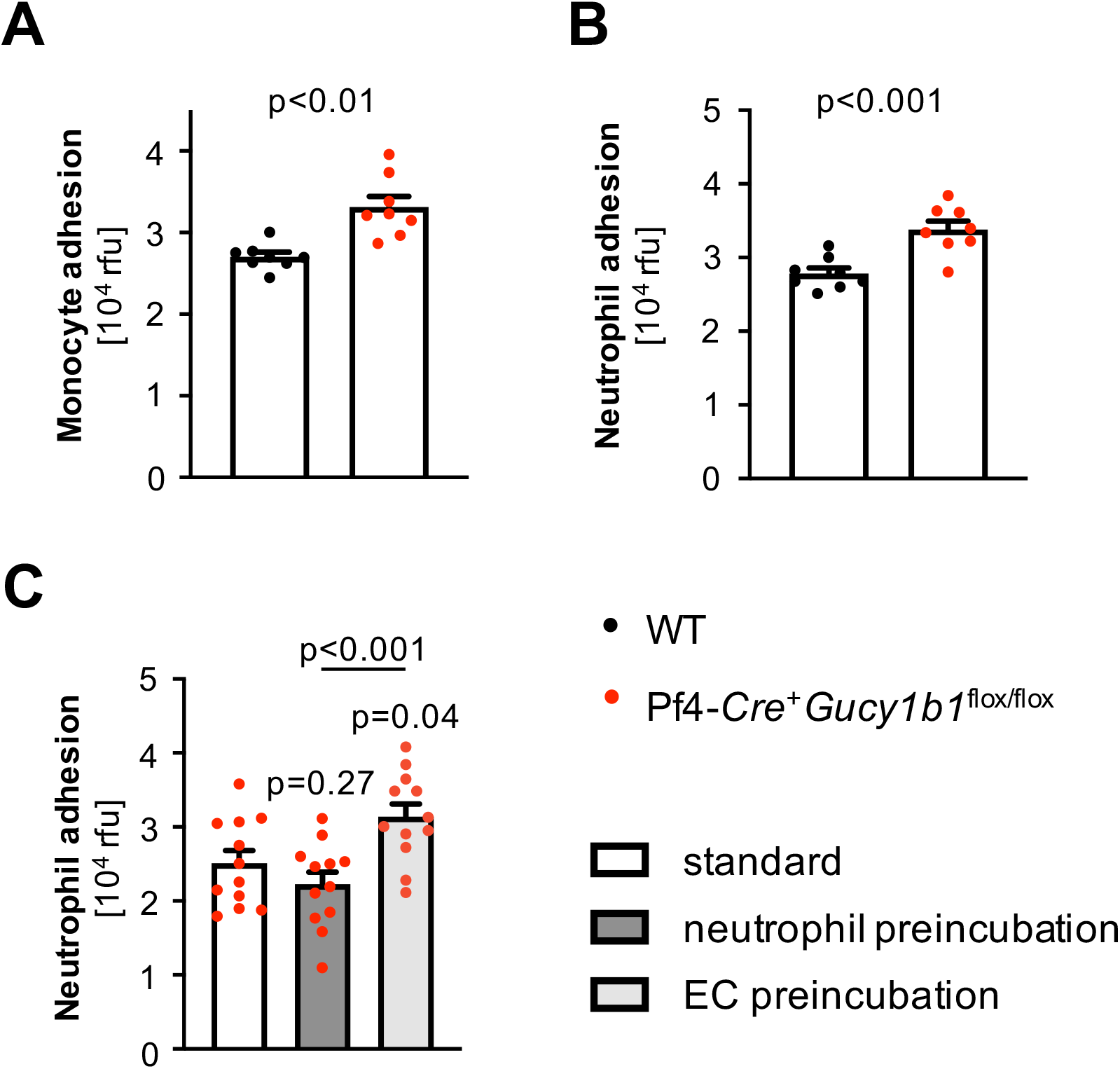
Influence of platelet sGC on leukocyte adhesion *in vitro*. **A, B**. WT monocyte (**A**) and neutrophil (**B**) adhesion to WT EC after incubation with supernatant of activated platelets isolated from either Pf4-*Cre*^+^*Gucy1b1*^flox/flox^ or WT mice. Each symbol represents one animal. Data are mean and s.e.m. Unpaired t-test. **C**. Quantification of neutrophil adhesion after preincubation of either EC or neutrophils with supernatant of activated platelets from Pf4-*Cre*^+^*Gucy1b1*^flox/flox^ mice in comparison to non-preincubation conditions. Each symbol represents one animal. Data are mean and s.e.m. RM one-way ANOVA with Tukey’s test for multiple testing. Abbreviations: *EC*, endothelial cells; *rfu*, relative fluorescence units.

### Reduced release of angiopoietin-1 by sGC-deficient platelets

We observed the influence of a soluble factor released by platelets on leukocyte adhesion to EC. To identify such factors, we next performed proteome profiling with supernatant of activated Pf4-*Cre*^+^*Gucy1b1*^flox/flox^ and WT platelets. Signal intensity analysis revealed lower angiopoietin-1 levels in the supernatant from Pf4-*Cre*^+^*Gucy1b1*^flox/flox^ platelets (3.2 ± 0.4 (n=7) vs. 6.7 ± 0.8 (n=8) arbitrary units, p<0.01; **Fig. 3A**). We next aimed at replicating this finding in an independent cohort of mice using an enzyme-linked immunosorbent assay (ELISA). Importantly, angiopoietin-1 levels were comparable in quiescent platelets (0.21 ± 0.01 vs. 0.23 ± 0.01 pg/10^3^ platelets, n=6 each, p=0.33; **Fig. 3B**) as well as in platelet poor plasma from Pf4-*Cre*^+^*Gucy1b1*^flox/flox^ and WT mice (0.6 ± 0.2 (n=6) vs. 0.4 ± 0.1 (n=5) ng/ml, p=0.46; **Fig. 3C**). In line with the explorative analysis displayed in **Fig. 3A**, we found Pf4-*Cre*^+^*Gucy1b1*^flox/flox^ platelets to release reduced amounts of angiopoietin-1 upon activation (30.4 ± 6.4 vs. 60.3 ± 4.7 ng/ml, n=6 each, p<0.01; **Fig. 3D**). Angiopoietin-1 is described to decrease particularly vascular endothelial growth factor-mediated adhesion of leukocytes to EC ^20^ and to bind to the Tie2 receptor on EC ^21^. We used the Tie2 inhibitor BAY-826 to investigate whether Tie2 inhibition influences leukocyte adhesion and found a 17% (± 1.2%, n=11 experiments, p=0.04) increase in leukocyte adhesion secondary to inhibiting the angiopoietin-1 receptor (**Fig. 3E**). This data demonstrate that angiopoietin-1 represents a candidate for mediating the effects of platelet sGC on leukocyte recruitment. Of note, inositol 1,4,5-trisphosphate receptor-associated cGMP kinase substrate (IRAG), which represents a downstream effector of cGMP specifically in modulating platelet activity, is encoded by the *IRAG1* ^22^ (previously IRAG or as human homologue *MRVI1*) gene and has also been associated with CAD by genome-wide association studies ^23^. To investigate whether differential angiopoietin-1 release is mediated via Irag1, we generated Pf4-*Cre*^+^*Irag1*^flox/flox^ mice and investigated platelet angiopoietin-1 release compared to WT mice. In contrast to Pf4-*Cre*^+^*Gucy1b1*^flox/flox^ platelets, we did not detect a difference between the genotypes in this experiment indicating that the influence of sGC on platelet angiopoietin-1 release is independent from Irag1 (**Suppl. Fig. S6**). It was previously shown that the genotype of the rs7692387 CAD risk allele at the *GUCY1A1* locus ^4^ influences *GUCY1A1* expression in different tissues ^8^ and sGC-α_1_ protein levels in platelets in particular ^7^. We therefore tested whether rs7692387 genotype is associated with angiopoetin-1 release from platelets in healthy human individuals (n=5 each, for characteristics see **Suppl. Table S2**) and found that homozygous carriers of the CAD risk allele G display lower angiopoietin-1 release compared to heterozygous or homozygous carriers of the non-risk allele (4.5 ± 0.7 vs. 8.3 ± 1.4 ng/ml, p=0.04; **Fig. 3F**).

**Figure 3:**
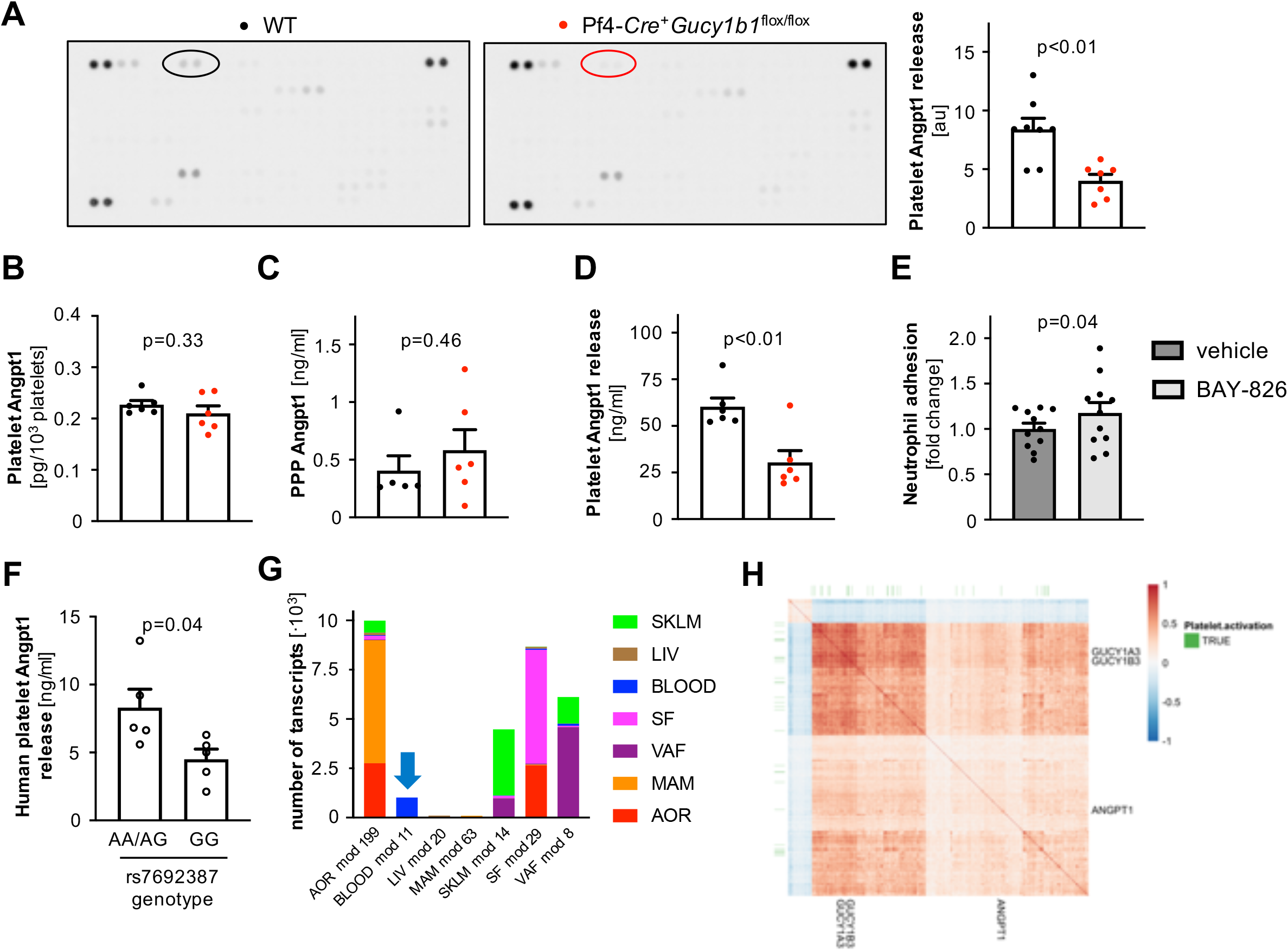
Platelet sGC influences release of angiopoietin-1. **A**. Left: Identification of angiopoietin-1 as differentially released protein from activated WT (black circle) and Pf4-*Cre*^+^*Gucy1b1*^flox/flox^ (red circle) platelets; right: quantification angiopoietin-1 signal in WT and Pf4-*Cre*^+^*Gucy1b1*^flox/flox^ mice. **B, C, D**. Quantification of platelet angiopoietin-1 content (**B**), platelet poor plasma angiopoietin-1 (**C**) and released angiopoietin-1 by ELISA (**D**). **E**. WT neutrophil adhesion to WT EC after incubation with supernatant of activated WT platelets in the absence and presence of the Tie2-inhibor BAY-826 (0.5 μM). **F**. Platelet angiopoietin-1 release in humans carrying the *GUCY1A1* (rs7692387) non-risk (AA, AG genotype) allele and homozygous carriers of the risk allele (GG genotype). Each symbol represents one animal or individual. Data are mean and s.e.m. Unpaired (**A-D, F**) or paired (**E**) t-test. **G**. STARNET co-expression modules containing *ANGPT1* from multitissue RNA-seq sampling of ∼600 CAD patients. The blue arrow denotes co-expression module 11 from whole blood samples (BLOOD). **H**. Heatmap of Pearson’s correlation coefficients of genes in co-expression module 11, showing positive correlation of *ANGPT1, GUCY1A3*, and *GUCY1B3* along with enrichment for platelet activation genes (FDR=6.863e-13, Enrichr, KEGG pathway). Abbreviations: *Angpt1*/*ANGPT1*, angiopoietin-1; *AOR*, aorta; *LIV*, liver; *MAM*, mammary artery; *PPP*, platelet poor plasma; *SF*, subcutaneous fat; *SKLM*, skeletal muscle; *VAF*, visceral fat.

To explore the role of angiopoietin-1 in relation to the genes encoding sGC in humans, we queried the STARNET database which contains bulk RNA-seq data from seven cardiometabolic tissues from patients undergoing coronary artery bypass graft surgery ^15^. *ANGPT1* was detected per tissue in seven distinct co-expression modules (**Fig. 3G**). Of note, *ANGPT1* was co-expressed with both *GUCY1A1* and *GUCY1B1* (encoding sGC-α_1_ and sGC-β_1_) in all tissues except mammary artery represented by co-expression module 63 (**Fig. 3G**). These data suggest ubiquitous presence and clinical variation in amounts of platelets. To investigate circulatory rather than tissue-resident platelets, we further analyzed co-expression module 11 from whole blood samples; it consisted of 1,016 genes and is estimated to account for 4.9% of CAD heritability by considering expression quantitative loci (eQTL) genes in a meta-analysis of nine GWAS using the restricted maximum likelihood method ^24^. Co-expression analysis revealed positive correlation between *GUCY1A1*/*GUCY1B1* and *ANGPT1* expression and enrichment for genes involved in the KEGG pathway platelet activation (**Fig. 3H**). Altogether this suggests that a substantial proportion of CAD heritability could be mediated by platelets. Furthermore, Gene Ontology enrichment analysis of this module (**Suppl. Table S2**) revealed, among others, “response to wounding” (p=4.71·10^−63^), “wound healing” (p=1.84·10^−60^), blood coagulation (p=1.99·10^−48^), and “regulation of locomotion” (p=6.64·10^−44^). These findings support the role of platelets and the interaction of angiopoietin-1 with sGC in human CAD.

### Modulation of angiopoietin-1 release and leukocyte recruitment by pharmacological sGC stimulation

The sGC represents a druggable target and sGC stimulators are used in different clinical scenarios, e.g. riociguat in pulmonary arterial hypertension and chronic thromboembolic pulmonary hypertension^25^ or vericiguat in chronic heart failure^26^. We next aimed at investigating whether modulation of the sGC using a vericiguat-like sGC stimulator (BAY-747) can influence the release of angiopoietin-1 and vascular inflammation. To this end, we first incubated platelets from WT mice with BAY-747 or vehicle and analyzed angiopoietin-1 release and leukocyte adhesion to endothelial cells. Stimulation of the sGC with BAY-747 doubled platelet angiopoietin-1 release (133.8 ± 6.5 vs. 64.0 ± 4.3 ng/ml, p<0.01; **Fig. 4A**) and reduced neutrophil adhesion to EC by 15.6% (± 6.5%, p=0.02; **Fig. 4B**). To investigate whether sGC stimulation *in vivo* is able to reduce recruitment from blood to the vascular wall, we fed *Ldlr*^-/-^ mice a Western diet containing 0 or 150 ppm BAY-747 and adoptively transferred GFP^+^ myeloid cells after six weeks: Here, we retrieved GFP^+^ monocytes admixed with neutrophils from naïve transgenic Ubc-GFP mice (all leukocytes express green fluorescent protein, GFP^+^) and injected the cells intravenously into wild-type mice (all cells are GFP^-^). After further 24 hours, the mice were sacrificed and numbers of GFP^+^ cells were determined using flow cytometry of aortic cell suspensions. The mice that received the Western diet containing the sGC simulator displayed reduced numbers of GFP^+^ cells (21.4 ± 3.3 vs. 42.8 ± 6.9 cells, n=8 each, p=0.01; **Fig. 4C**). These data indicate that pharmacological sGC stimulation is able to modulate platelet angiopoietin-1 release, *in vitro* leukocyte adhesion and *in vivo* leukocyte recruitment.

**Figure 4:**
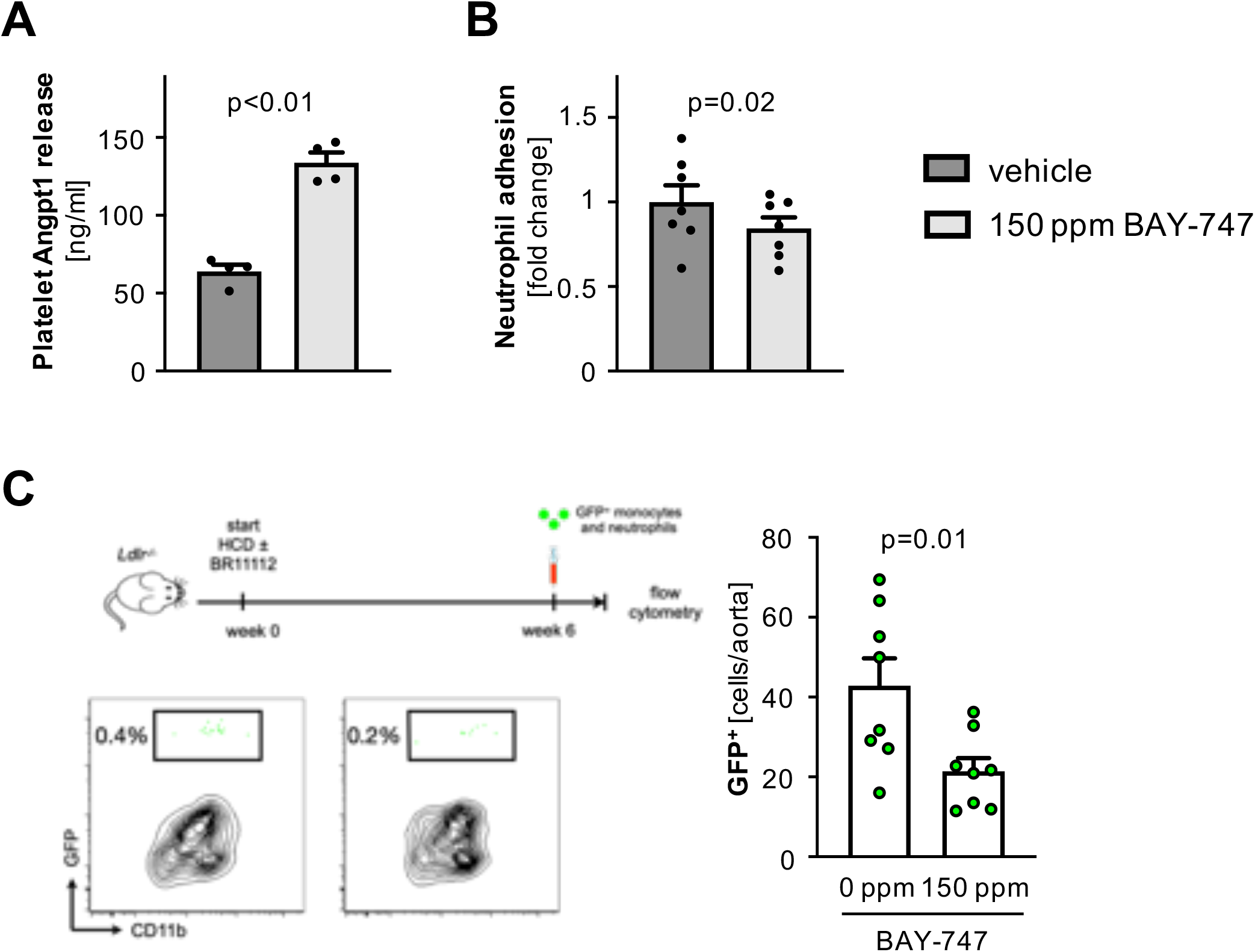
Modulation of angiopoietin-1 release and leukocyte recruitment by pharmacological sGC stimulation. **A**. Angiopoietin-1 release by WT platelets incubated with 150 ppm BAY-747 or vehicle. **B**. WT neutrophil adhesion to WT EC after incubation with supernatant from activated WT platelets that were preincubated with either vehicle or 150 ppm BAY-747. Each symbol represents one animal. Data are mean and s.e.m. Paired t-test. **C**. Adoptive transfer of GFP^+^ leukocytes: study scheme, flow cytometry plot and (left) and quantification (right) of GFP^+^ cells. Each symbol represents one animal. Unpaired t-test.

### Therapeutic potential of stimulating sGC in atherosclerosis

The findings displayed in **Fig. 4** suggest a potential benefit from sGC stimulation under proatherogenic condition. To determine whether such treatment is able to reduce atherosclerotic plaque formation, we again fed atherosclerosis-prone *Ldlr*^-/-^ mice a Western diet for ten weeks. In one group, the diet contained 150 ppm BAY-747 (treatment group) while the other group received a western diet without (=0 ppm) BAY-747 (control group). Under steady state conditions, BAY-747 plasma levels in the male and female mice receiving BAY-747 were 61.7 ± 1.4 μg/l and 40.3 ± 1.7 μg/l (n=6, each), respectively; in the control group, BAY-747 was not detectable in plasma as expected (**Suppl. Fig. S7**). Both groups had elevated serum cholesterol levels without significant difference between the two groups; similarly, platelet count, blood leukocytes, and body weight after diet were comparable (**Supp. Fig. S8**). In the aortic root, we found significantly reduced atherosclerotic plaque formation in mice of the treatment group (62.5 ± 16.1 (n=7) vs. 123.1 ± 18.6 (n=9) μm^2^, p=0.03; **Fig. 5A**). We furthermore detected fewer leukocytes in aortic roots of those animals (30.3 ± 5.8 (n=8) vs. 50.5 ± 6.3 (n=11) % of plaque area, p=0.04; **Fig. 5B**). In cell suspensions of the whole aorta, we furthermore found less numerous leukocytes (23,546 ± 2,235 vs. 37,984 ± 3,607 Cd11b^+^ cells/aorta, n=12 each, p<0.01), macrophages (13,915 ± 1,550 vs. 22,156 ± 2,737 macrophages/aorta, n=12 each, p=0.02; **Fig. 5C**), Ly6C^high^ monocytes (1,485 ± 348 vs. 2,688 ± 531 Ly6C^high^ monocytes/aorta, n=12 each, p=0.07; **Fig. 5C**), and neutrophils (1,201 ± 192 (n=11) vs. 1,885 ± 232 (n=12) neutrophils/aorta, p=0.04; **Fig. 5C**) in mice treated with the sGC stimulator compared to mice receiving the control diet.

**Figure 5:**
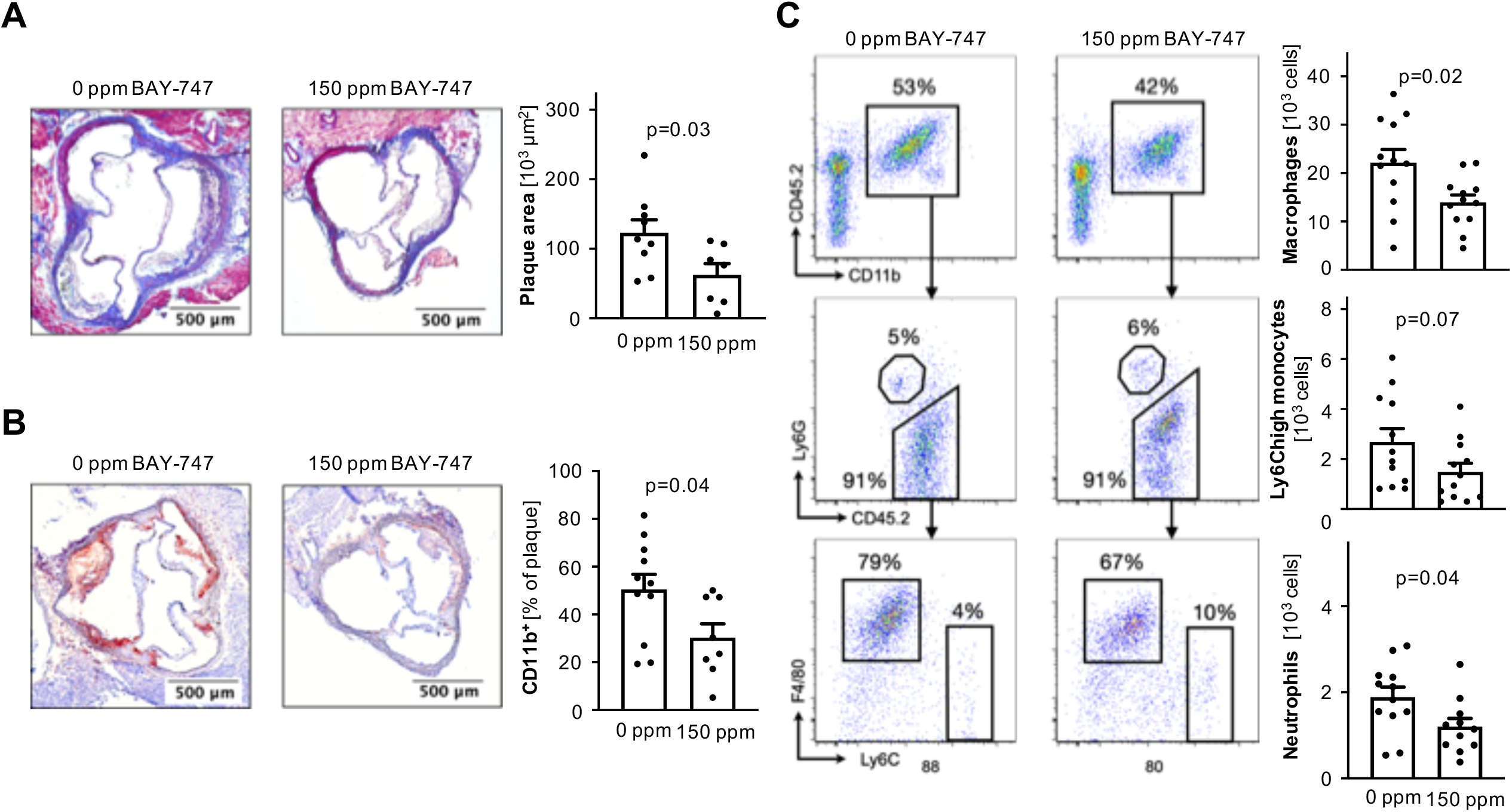
Pharmacological sGC stimulation influences atherosclerotic plaque formation and vascular inflammation. **A**. Aortic root atherosclerotic plaques in *Ldlr*^-/-^ mice which were fed a western diet for ten weeks containing 0 (control group) or 150 ppm BAY-747. **B**. Cd11b^+^ area of aortic roots in mice from the control and treatment group. **C**. Quantification of vascular inflammation by flow cytometry analysis of aortic cell suspensions of mice in the control and treatment group. Each symbol represents one mouse. Data are mean and s.e.m. Unpaired t-test. One outlier was removed in panel C in the analysis of neutrophils according to the ROUT method.

Blood leukocyte numbers as determined by flow cytometry and leukocyte subsets were comparable between both groups (**Suppl. Fig. S9**). Taken together and in conjunction with the data presented in **Fig. 4C**, these data indicate that pharmacological sGC stimulation reduces atherosclerotic plaque formation and vascular inflammation via dampened recruitment of inflammatory leukocytes from blood to plaque.

## Discussion

NO-sGC-cGMP-signaling has important functions in a variety of cell types. For instance, increasing intracellular cGMP levels inhibits migration of vascular smooth muscle cells and aggregation of platelets, respectively ^27,28^. The observation that genes encoding key proteins in this pathway were associated with CAD and premature MI by genome-wide association ^4,23,29,30^ and exome sequencing studies ^5,8,31^ renders an important role in the pathophysiology of coronary atherosclerosis likely. In this study, we sought to specifically investigate the role of platelet sGC in atherosclerosis because we found that carriers of the common, non-coding risk variant rs7692387 identified by genome-wide association studies ^4^ displayed reduced expression of sGC in platelets and – as a consequence – reduced inhibition of platelet aggregation upon NO stimulation ^7^. In addition, we observed that in contrast to the general population, individuals that are homozygous for this variant had a benefit from aspirin treatment in primary prevention of cardiovascular events ^10^. In a series of *in vivo* and *in vitro* experiments, we here observed larger atherosclerotic plaques in aortic roots of mice lacking platelet sGC compared to sGC wildtype mice. Platelet sGC has just recently been described to act as an endogenous brake on platelet aggregation ^32^. Indeed, it has been hypothesized that platelets adhering to endothelial or plaque erosions may be activated more or less depending of the availability of sGC. This may stimulate inflammation locally and, subsequently, facilitate atherosclerotic plaque formation. While this remains a hypothesis, we here report evidence that platelet sGC influences the release of the soluble factor angiopoietin-1 with reduced amounts released by platelets lacking sGC. The role of angiopoietin-1 in atherosclerosis is controversial as there are reports describing a protective ^20,21,33^ as well as a deleterious role ^34^. However, given the cellular findings describing the inhibitory effect of angiopoietin-1 on leukocyte adhesion and vascular atherosclerosis reported in the literature ^20^ – which were confirmed in this study – and the beneficial role of angiopoietin-1-Tie2-signaling in inflammatory diseases in general ^35^, it represents an interesting candidate mediating downstream effects of platelet sGC signaling. Importantly, the role of sGC in platelets seems to exceed the role as a brake on aggregation: while sGC passivates platelet aggregation, it enhances the release of angiopoietin-1. The notion that this effect is independent of aggregation is further supported by the findings i) that Irag1, the mediator of cGMP-dependent inhibition of platelet aggregation ^36^ was not involved in angiopoietin-1 release and ii) that angiopoietin-1 release was induced by shaking but not, e.g., ADP or arachidonic acid. It is important to acknowledge that the content of angiopoietin-1 in WT and KO platelets was similar further indicating a direct link between sGC availability and angiopoietin-1 release. This is further emphasized by the finding that also a genetically determined reduction – but not lack of – sGC in platelets of carriers of the CAD-associated risk variant rs7692387 was associated with a reduction in angiopoietin-1 release. This, on one hand, supports the translational relevance of this by nature artificial *in vitro* and animal studies; on the other hand, it raises the question whether modulators of sGC function could be used to prevent and treat coronary atherosclerosis. Indeed, stimulators of the sGC are emerging as therapeutic compounds for different cardiovascular diseases (for an overview see ^37^). Recently, the sGC stimulator vericiguat was found to lower the risk of death from cardiovascular causes or hospitalization for heart failure in patients with heart failure with reduced ejection fraction ^26^. Data on atherosclerotic plaque formation and ischemic cardiovascular events are however, so far lacking.

To this end, we investigated whether sGC stimulation using a vericiguat-like stimulator, BAY-747, is able to modulate the reported cellular and phenotypic effects. First, we found that sGC stimulation is able to increase angiopoietin-1 release and subsequently reduce leukocyte adhesion to endothelial cells *in vitro*. As we postulate that sGC contributes to reducing vascular inflammation, we next studied whether sGC stimulation is able to alter the recruitment of inflammatory leukocytes from blood to plaque. In an adoptive transfer experiment, we observed a reduction in leukocyte recruitment which is furthermore in line with previous reports that showed anti-inflammatory effects of cGMP-increasing pharmacological compounds ^38,39^. The finding of multiple genome-wide significant hits in genes which encode proteins that are tightly involved in the formation (*NOS3* ^29^, *GUCY1A1* ^4^), the fate (*PDE5A* ^30,31^), or mediating the downstream effects of cGMP (*IRAG1* ^23^, *PDE3A* ^40^) and the observation that sGC levels are reduced in atherosclerotic tissues ^41^ increase the likelihood that targeting sGC might be beneficial in CAD. In atherosclerosis-prone mice on a Western diet, stimulating sGC with BAY-747 led to a reduction in atherosclerotic plaque formation and vascular inflammation warranting further investigation into this promising pharmacological treatment strategy. Further evidence might be taken from a recent study which compared two pharmacological strategies which are used to treat erectile dysfunction: compared to alprostadil, i.e. prostaglandin E1, treatment with the inhibitor of phosphodiesterase 5A, sildenafil, was associated with a reduced risk of all-cause mortality and myocardial infarction in men suffering from CAD ^42^. To definitively prove a beneficial role in CAD, however, prospective clinical trial data are needed.

Taken together, we here provide further evidence for a crucial role of platelets in atherosclerosis in general – and of platelet sGC in particular. As shown by our *in vitro* studies using human and murine biospecimen and *in vivo* studies, we postulate an endogenous inhibitory role of platelet sGC on EC-mediated leukocyte recruitment (**Fig. 6**). We are aware that platelets are not the only cell types in which sGC activity and genetic variants modulating its availability influence CAD risk. Modulating sGC activity, especially using stimulators, might nevertheless be a promising therapeutic strategy exceeding the effects on platelet sGC.

**Figure 6:**
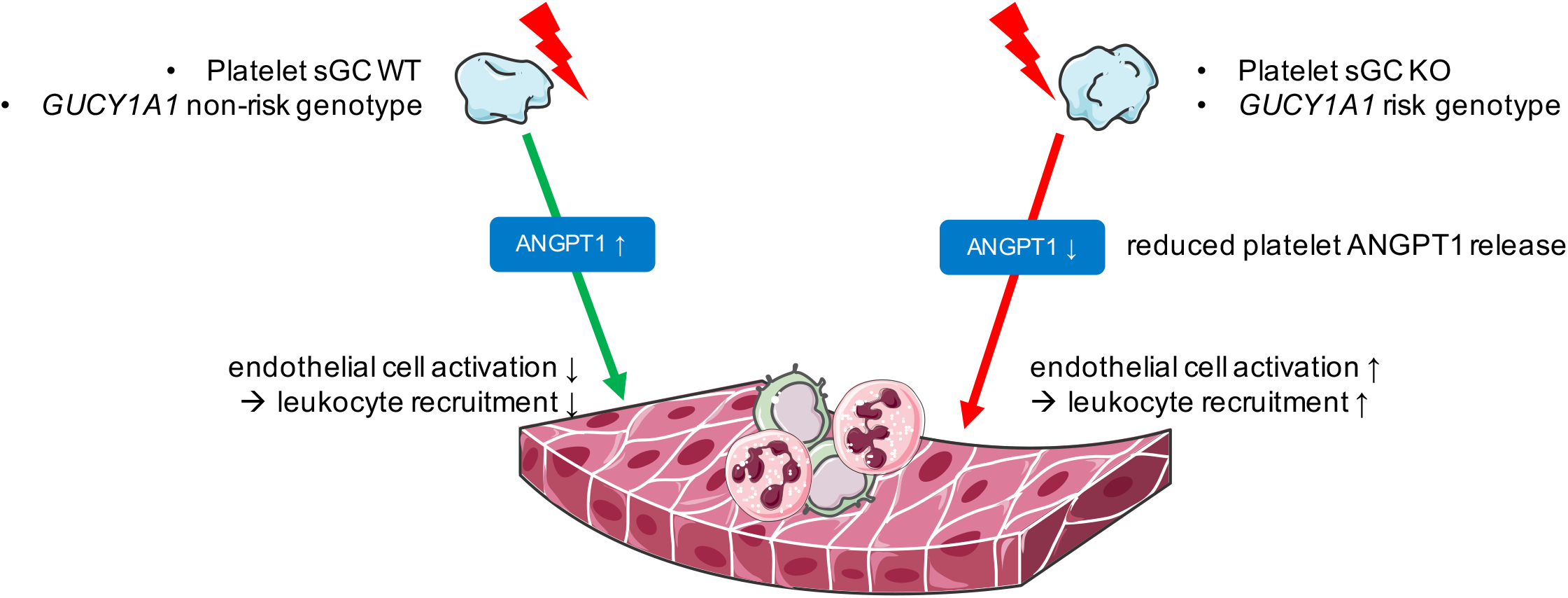
Graphical Abstract. In the event of platelet activation, e.g. by shear stress, sGC counterbalances proinflammatory activation of EC, e.g., by release of angiopoietin-1 (ANGPT1). If sGC levels are reduced – e.g. in the mouse model used in this study or in platelets of homozygous carriers of the CAD-associated risk variant – less ANGPT1 is released. Subsequently enhanced EC activation and leukocyte recruitment contribute to atherosclerotic plaque formation.

Our study has several limitations. First, this is an *in vitro* and mouse *in vivo* study that cannot resemble the human physiology and pathophysiology. However, the finding that lack of platelet sGC in mice and genetically determined reduced platelet sGC-α_1_ in humans both reduce release of platelet angiopoietin-1 supports a possible translation. Furthermore, similar to humans a genetically determined reduction in *Gucy1a1* mRNA was shown to be associated with increased atherosclerotic plaque formation in the hybrid mouse diversity panel ^7,43^. Second, although we have shown that reduced angiopoietin-1 release and enhanced leukocyte adhesion to EC is linked to reduced or lacking platelet sGC availability and that stimulation of the sGC is able to modulate these downstream effects, the exact molecular mechanism linking sGC and angiopoietin-1 release remains to be explored. While it is well known that platelets contain at least three types of secretory granules, with α- granules that also harbor angiopoietin-1 ^44^ being the most abundant type, there is also evidence for the existence of functionally distinct subpopulations within α-granules ^45^, which may allow selective release of their contents by different stimuli ^46,47^. A similar observation has been reported in neutrophils regarding storage of angiopoietin-1 and VEGF ^48^. We further provide evidence that it is independent from IRAG1. Third, we know that angiopoietin-1 is likely not the only mediator of sGC effects on atherosclerosis. Lastly, we cannot generalize a benefit of sGC stimulation to humans. Our study can yet, in addition to its biological implications, be regarded as hypothesis-generating for future clinical trials investigating whether modulating cGMP formation is useful in addition to reduce, e.g., LDL-cholesterol levels and residual inflammatory risk.

## Acknowledgments

The authors thank Christopher Wolf for technical assistance. Figure 6 contains modified image material available at Servier Medical Art under a Creative Commons Attribution 3.0 Unported License. The authors’ work is funded by the Corona Foundation as part of the Junior Research Group Translational Cardiovascular Genomics (S199/10070/2017, to T.K.) and the German Research Foundation (DFG) as part of the collaborative research centers SFB 1123 (B02, to T.K. and H.S.), TRR 267 (B06, to H.S.), and STRESS_638675 (to H.B.S.). Further grants were received from the European Research Council (ERC) under the European Union’s Horizon 2020 research and innovation program (grant agreement No 759272, to H.B.S.) and the German Heart Foundation (Deutsche Herzstiftung; F/28/17, to H.B.S.) as well as the German Federal Ministry of Education and Research (BMBF) within the framework of ERA-NET on Cardiovascular Disease (ERA-CVD: grant JTC2017_21-040, to H.S. and J.L.M.B.) and within the scheme of target validation (BlockCAD: 6GW0198K, to H.S.). Additional support was received from the British Heart Foundation/DZHK collaborative project “Genetic discovery-based targeting of the vascular interface in atherosclerosis”.

## One-sentence summary

Loss of platelet soluble guanylyl cyclase activity contributed to atherosclerotic plaque formation and vascular inflammation while pharmacological stimulation of the enzyme represented a promising counteracting strategy.

