## Supplemental Material for "Loss of soluble guanylyl cyclase in platelets contributes to atherosclerotic plaque formation and vascular inflammation"

^#^	Heribert Schunkert and Thorsten Kessler provided equal supervision of this work.

Corresponding authors:

Hendrik B Sager, MD, Heribert Schunkert, MD & Thorsten Kessler, MD; German Heart Centre Munich; Department of Cardiology; Technical University of Munich; Lazarettstr. 36 · 80636 Munich, Germany; Phone: +49 89 1218 4073 · Fax: +49 89 1218 4013;

*Protein extraction*

Lung and aorta were harvested from Pf4-*Cre*^+^*Gucy1b1*^+/flox^ and Pf4-*Cre^+^Gucy1b1*^flox/flox^ mice after perfusion of organs with PBS (Sigma Aldrich, St. Louis, MA) and snap-frozen in liquid nitrogen. For protein isolation, specimens were placed in ice-cold radioimmunoprecipitation assay buffer (RIPA, Cell Signaling Technology, Danvers, MA) supplemented with 1:100 protease inhibitor cocktail (1861278, Thermo Fisher, Waltham, MA) and disrupted using an electric tissue homogenizer (Omni International, Kennesaw, GA) on ice. For generation of thrombocyte lysates, platelets were collected from heparinized full blood as stated before and resuspended in RIPA buffer supplemented with protease inhibitor to a number of 2 x 10^8^ cells/ml. Cells were disrupted by intermittent sonication three times for 30 seconds in an ice-bath. Protein concentrations were determined using a bicinchoninic acid (BCA) assay (23227, Thermo Fisher) according to the manufacturer’s protocol.

*Immunoblotting*

Samples were supplemented with 4X Laemmli buffer containing 355 mM 2-mercaptoethanol (Bio-Rad Laboratories, Hercules, CA) and denatured for 5 min at 95°C. Proteins were separated by SDS-gel electrophoresis using 4-20 % Mini-PROTEAN® TGX™ precast gradient gels (Bio-Rad, Hercules, CA) at 100 V for one hour in 1X Tris/glycine/SDS buffer (Bio-Rad). Wet blotting (25 mM Tris, 192 mM glycine, 20 % v/v methanol, pH 8.3) was performed at 100 V for 90 minutes using methanol-activated polyvinylidene difluoride (PVDF) membranes (Merck Millipore, Darmstadt, Germany). Membranes were blocked for one hour at room temperature in PBS-T (PBS with 0.2 % Tween, Sigma Aldrich) containing 5 % nonfat dry milk powder (NFDM, AppliChem, Darmstadt, Germany). Primary antibodies (anti-mβ1, directed against the β_1_-subunit of the sGC ^1^ diluted 1:1,000 and GAPDH, 8884S, Cell Signaling Technology, 1:10,000 in 2.5% NFDM-PBS-T) were incubated at 4 °C overnight, followed by a one-hour incubation at room temperature with anti-rabbit HRP-conjugated secondary antibody (7074, Cell Signaling Technology, 1:100,000 in 2.5% NFDM-PBS-T). For signal detection, membranes were developed with SuperSignal™ West Dura Extended Duration Substrate (Thermo Fisher) according to the manufacturer´s recommendations and signal intensities were detected using an ImageQuant LAS 400 imaging system (GE Healthcare Life Sciences, Freiburg, Germany).

*Platelet aggregation*

Platelet rich plasma (PRP) was collected from Pf4-*Cre*^+^*Gucy1b1*^+/flox^ and Pf4-*Cre^+^Gucy1b1*^flox/flox^ mice as stated in the manuscript and thrombocyte count was measured using an automated hematology analyzer (Sysmex Corp, Kobe, Japan). PRP was centrifuged at 700 g for 10 minutes to receive platelet poor plasma used for blanking. Samples were incubated at 37°C in glass cuvettes with constant stirring on an 8-channel personal computer-controlled platelet aggregation profiler (PAP-8, Biodata Corp, Horsham, PA) and thrombocyte aggregation was induced by addition of ADP (final concentration 2 µmol/l; möLaboratory GmbH, Langenfeld, Germany). Platelet aggregation was recorded over five minutes to measure area under the curve (AUC). In the next step, samples were stimulated with sodium nitroprusside (final concentration 10 µmol/l, Sigma Aldrich) prior to the addition of ADP for two minutes and platelet aggregation measured as stated above.

*Determination of BAY-747 serum/plasma concentration*

BAY 747 exposure was quantified in plasma using an LC-system for mass separation (Kinetex 5 µm C18 100 A LC Column 150*4.6 mm) coupled to a 4500 Triple Quad Sciex mass analyzer (positive mode). A generic internal standard was added to the samples. A 5 point calibration curve and quality control samples were used for relative quantification. Plasma were obtained from 6 mice per group. Data are mean + SEM.

**Supplemental Figures**

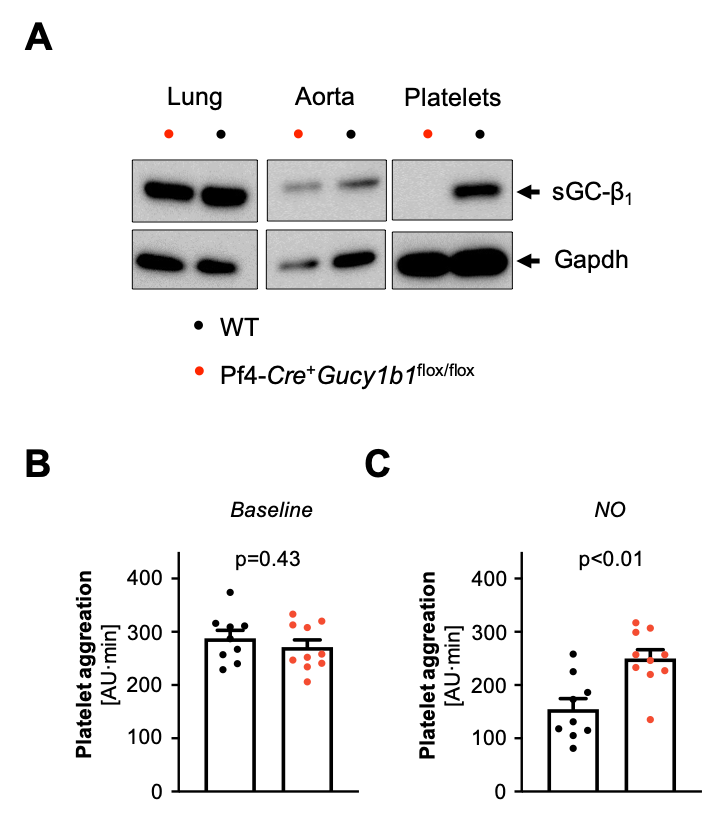

**Suppl. Figure S1**: Pf4-*Cre*^+^*Gucy1b1*^flox^/^flox^ compared to WT mice. **A**. Expression of sGC-β1 in lung, aorta, and platelets. **B, C**. Platelet aggregation after adenosine diphosphate (ADP) stimulation under baseline conditions (**B**) and secondary to stimulation with the nitric oxide (NO) donor sodium nitroprusside (**C**). Each symbol represents one animal. Data are mean and s.e.m. Unpaired t-test.

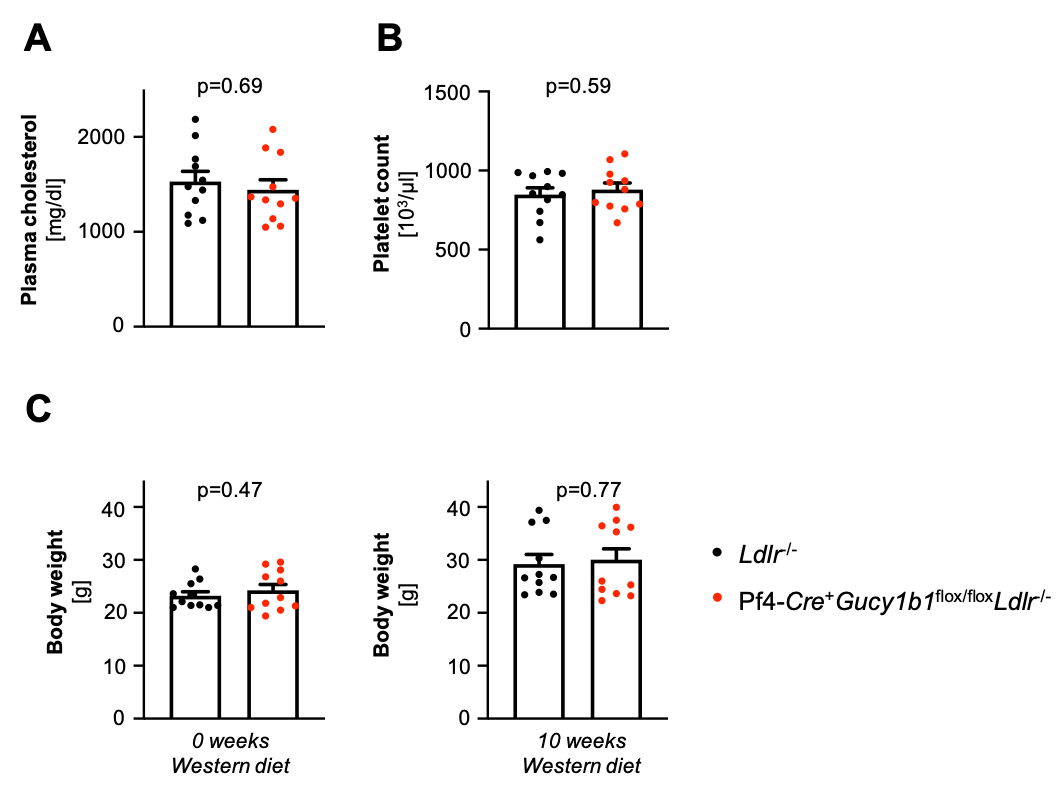

**Suppl. Figure S2**: Pf4-*Cre*^+^*Gucy1b1*^flox^/^flox^*Ldlr*^-/-^ compared to *Ldlr*^-/-^ mice after Western diet (see **Fig. 1E**). **A**. Serum cholesterol levels. **B**. Platelet count. **C**. Blood leukocytes. **D**. Body weight at baseline (left) and after ten weeks (right). Each symbol represents one animal. Data are mean and s.e.m. Unpaired t-test.

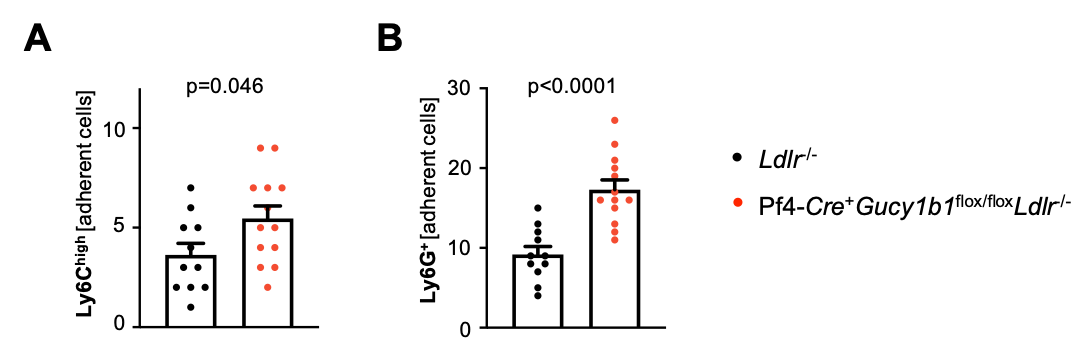

**Suppl. Figure S3**: Leukocyte adhesion to atherosclerotic plaques of Pf4-*Cre*^+^*Gucy1b1*^flox^/^flox^*Ldlr*^-/-^ compared to *Ldlr*^-/-^ mice after Western diet assessed by fluorescence intravital microscopy. **A**. Ly6C^high^ monocytes. **B**. Neutrophils. Each symbol represents one animal. Data are mean and s.e.m. Unpaired t-test.

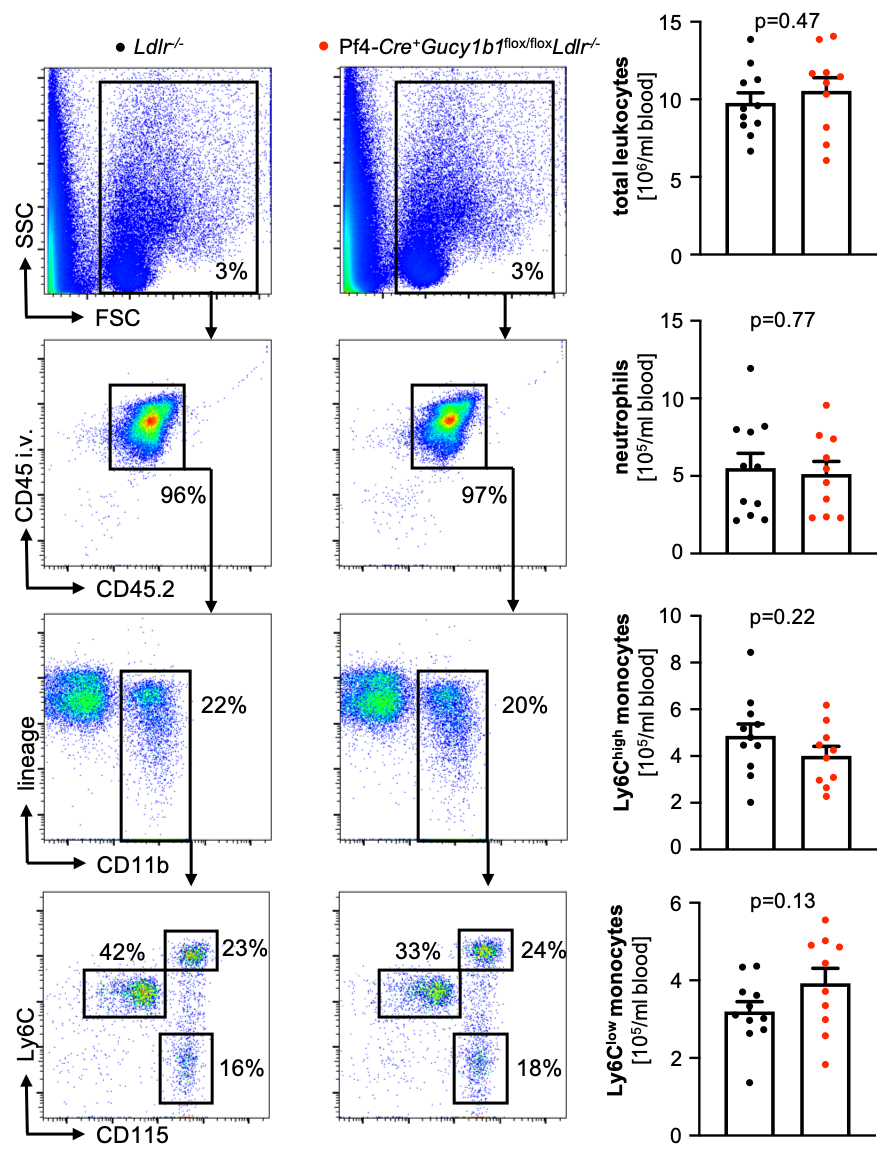

**Suppl. Figure S4**: Blood leukocyte numbers and subsets in Pf4-*Cre*^+^*Gucy1b1*^flox^/^flox^*Ldlr*^-/-^ compared to *Ldlr*^-/-^ mice after Western diet (see **Fig. 1E**). Each symbol represents one animal. Data are mean and s.e.m. Unpaired t-test. One outlier was removed in the analysis of Ly6C^low^ monocytes according to the ROUT method.

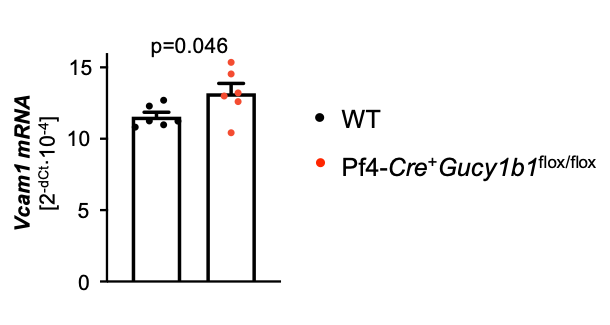

**Suppl. Figure S5**: *Vcam1* expression of wildtype (WT) endothelial cells after incubation with supernatant of activated WT or Pf4-*Cre*^+^*Gucy1b1*^flox^/^flox^ platelets. Each symbol represents one animal. Data are mean and s.e.m. Unpaired t-test.

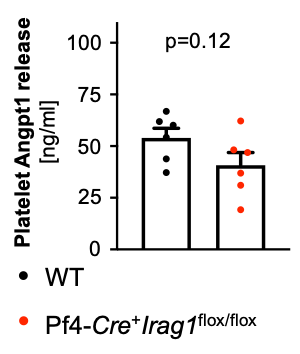

**Suppl. Figure S6**: Angiopoietin-1 (Angpt1) release by wild type (WT) or Pf4-*Cre*^+^*Irag1*^flox^/^flox^ platelets after activation by shaking. Each symbol represents one animal. Data are mean and s.e.m. Unpaired t-test.

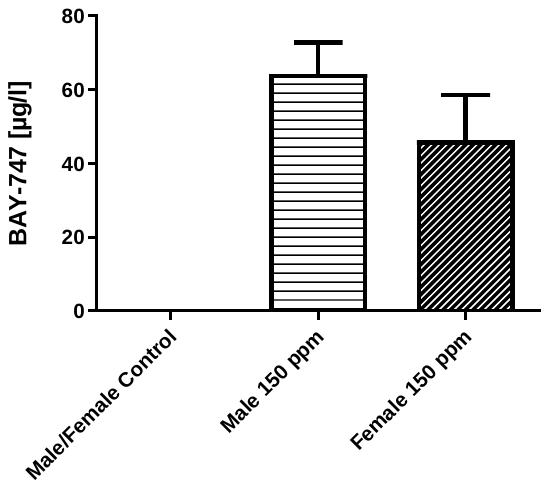

**Suppl. Figure S7**: Plasma concentrations of the soluble guanylyl cyclase stimulator BAY-747 in mice receiving a Western diet containing 0 ppm or 150 ppm BAY-747. Each symbol represents one animal. Data are mean and s.e.m. N=6 mice per group and gender.

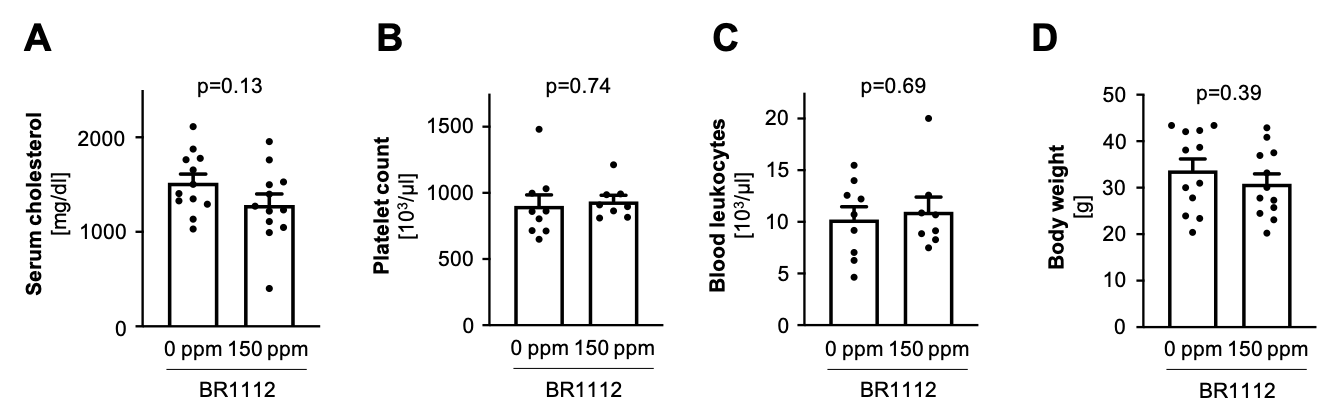

**Suppl. Figure S8**: *Ldlr*^-/-^ mice receiving 0 ppm compared to *Ldlr*^-/-^ mice receiving 150 ppm BAY-747 after ten weeks of Western diet. **A**. Serum cholesterol levels. **B**. Platelet count. **C**. Blood leukocytes. **D**. Body weight after. Each symbol represents one animal. Data are mean and s.e.m. Unpaired t-test.

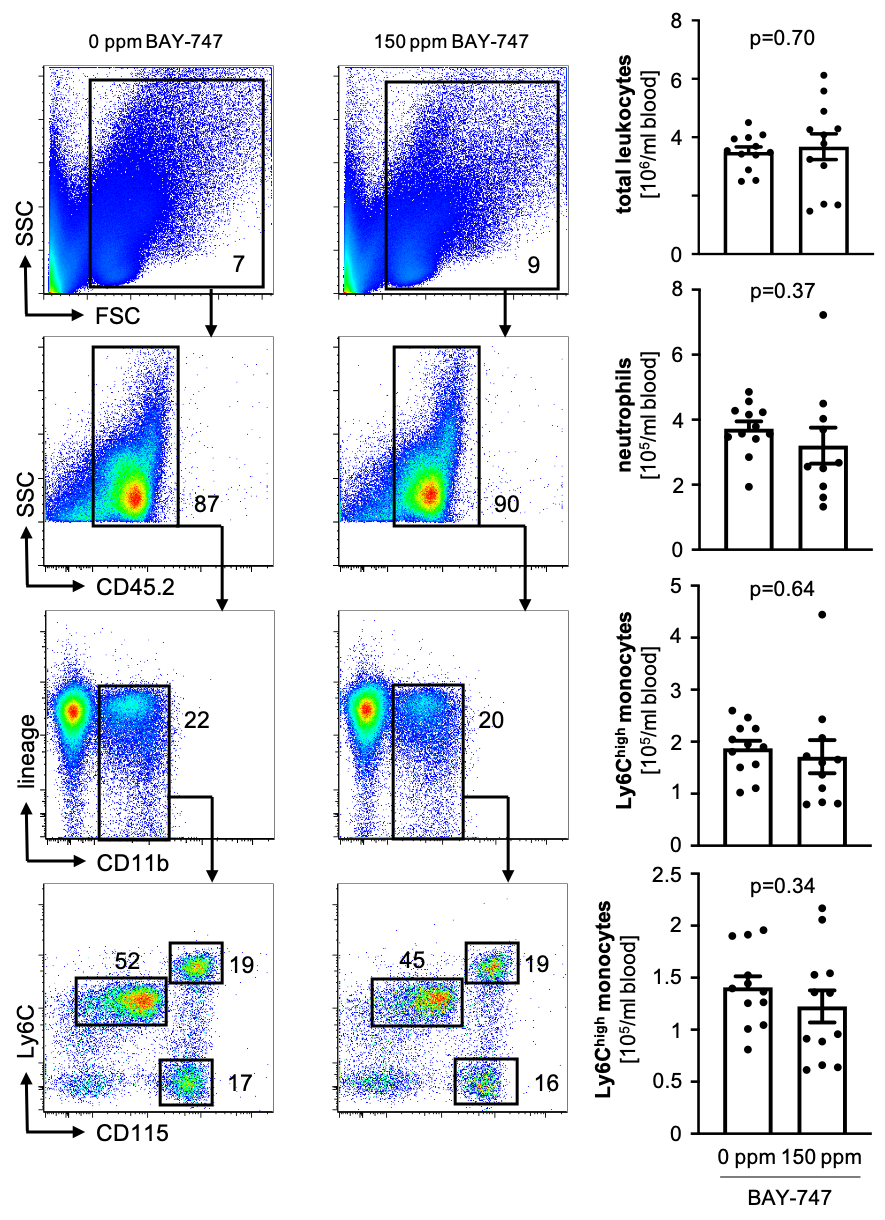

**Suppl. Figure S9**: Blood leukocyte numbers and subsets in *Ldlr*^-/-^ mice receiving 0 ppm compared to *Ldlr*^-/-^ mice receiving 150 ppm BAY-747 after ten weeks of Western diet. Each symbol represents one animal. Data are mean and s.e.m. Unpaired t-test.

**Supplemental Tables**

**Suppl. Table S1**: Distribution of data and statistical test used. *, Kolmogorov-Smirnov test.

| **Figure, panel** | **Comparison** | **Normal distribution* (p-value)** | **Test used** |
| --- | --- | --- | --- |
| 1, A | Plaque size; *Ldlr*^-/-^ vs. Pf4-*Cre*^+^*Gucy1b1*^flox/flox^*Ldlr*^-/-^ | Yes (>0.10) | Unpaired t-test |
| 1, B | Plaque size aorta *en face*; *Ldlr*^-/-^ vs. Pf4-*Cre*^+^*Gucy1b1*^flox/flox^*Ldlr*^-/-^ | Yes (>0.10) | Unpaired t-test |
| 1, C | MOMA-2+ area; *Ldlr*^-/-^ vs. Pf4-*Cre*^+^*Gucy1b1*^flox/flox^*Ldlr*^-/-^ | Yes (>0.10) | Unpaired t-test |
| 1, D | Adherent leukocytes; *Ldlr*^-/-^ vs. Pf4-*Cre*^+^*Gucy1b1*^flox/flox^*Ldlr*^-/-^ | Yes (>0.10) | Unpaired t-test |
| 1, E | Macrophages, FACS; *Ldlr*^-/-^ vs. Pf4-*Cre*^+^*Gucy1b1*^flox/flox^*Ldlr*^-/-^ | No (=0.03) | Mann-Whitney test |
|  | Ly6Chigh monocytes, FACS; *Ldlr*^-/-^ vs. Pf4-*Cre*^+^*Gucy1b1*^flox/flox^*Ldlr*^-/-^ | Yes (>0.10) |  |
|  | Ly6G+ leukocytes, FACS; *Ldlr*^-/-^ vs. Pf4-*Cre*^+^*Gucy1b1*^flox/flox^*Ldlr*^-/-^ | Yes (>0.10) |  |
| 2, A | *In vitro* monocyte adhesion; WT vs. Pf4-*Cre*^+^*Gucy1b1*^flox/flox^ | Yes (>0.10) | Paired t-test |
| 2, B | *In vitro* neutrophil adhesion; WT vs. Pf4-*Cre*^+^*Gucy1b1*^flox/flox^ | Yes (>0.10) | Paired t-test |
| 2, C | *In vitro* neutrophil adhesion; Pf4-*Cre*^+^*Gucy1b1*^flox/flox^; standard vs. neutrophil vs. EC preincubation | Yes (>0.10) | RM one-way ANOVA, Tukey multiple testing |
| 3, A | Platelet Angpt1 release, proteome profiler; WT vs. Pf4-*Cre*^+^*Gucy1b1*^flox/flox^ | Yes (>0.10) | Unpaired t-test |
| 3, B | Platelet Angpt1 content, ELISA; WT vs. Pf4-*Cre*^+^*Gucy1b1*^flox/flox^ | Yes (>0.10) | Unpaired t-test |
| 3, C | Platelet poor plasma Angpt1, ELISA; WT vs. Pf4-*Cre*^+^*Gucy1b1*^flox/flox^ | Yes (>0.10) | Unpaired t-test |
| 3, D | Platelet Angpt1 release, ELISA; WT vs. Pf4-*Cre*^+^*Gucy1b1*^flox/flox^ | Yes (>0.10) | Unpaired t-test |
| 3, E | *In vitro* neutrophil adhesion; WT, DMSO vs. BAY-826 | Yes (>0.10) | Paired t-test |
| 3, F | Human platelet ANGPT1 release, ELISA; rs7692387 AA/AG vs. GG genotype carriers | Yes (>0.10) | Unpaired t-test |
| 4, A | Platelet Angpt1 release, ELISA, WT; vehicle vs. BAY-747 | Yes (>0.10) | Paired t-test |
| 4, B | *In vitro* neutrophil adhesion, WT; vehicle vs. 150 ppm BAY-747 | Yes (>0.10) | Paired t-test |
| 4, C | GFP+ leukocytes, FACS; *Ldlr*^-/-^; 0 vs. 150 ppm BAY-747 | Yes (>0.10) | Unpaired t-test |
| 5, A | Plaque size; *Ldlr*^-/-^; 0 vs. 150 ppm BAY-747 | Yes (>0.10) | Unpaired t-test |
| 5, B | Cd11b+ area; *Ldlr*^-/-^; 0 vs. 150 ppm BAY-747 | Yes (>0.10) | Unpaired t-test |
| 5, C | Cd11b+ leukocytes, FACS; *Ldlr*^-/-^; 0 vs. 150 ppm BAY-747 | Yes (>0.10) | Unpaired t-test |
|  | Macrophages, FACS; *Ldlr*^-/-^; 0 vs. 150 ppm BAY-747 | Yes (>0.10) |  |
|  | Ly6Chigh monocytes, FACS; *Ldlr*^-/-^; 0 vs. 150 ppm BAY-747 | Yes (>0.10) |  |
|  | Ly6G+ leukocytes, FACS; *Ldlr*^-/-^; 0 vs. 150 ppm BAY-747 | Yes (>0.10) |  |
| S1, B | Platelet aggregation, baseline; WT vs. Pf4-*Cre*^+^*Gucy1b1*^flox/flox^ | Yes (>0.10) | Unpaired t-test |
| S1, C | Platelet aggregation, after NO; WT vs. Pf4-*Cre*^+^*Gucy1b1*^flox/flox^ | Yes (>0.10) | Unpaired t-test |
| S2, A | Serum cholesterol, after western diet; *Ldlr*^-/-^ vs. Pf4-*Cre*^+^*Gucy1b1*^flox/flox^*Ldlr*^-/-^ | Yes (>0.10) | Unpaired t-test |
| S2, B | Platelets, after western diet; *Ldlr*^-/-^ vs. Pf4-*Cre*^+^*Gucy1b1*^flox/flox^*Ldlr*^-/-^ | Yes (>0.10) | Unpaired t-test |
| S2, C | Leukocytes, after western diet; *Ldlr*^-/-^ vs. Pf4-*Cre*^+^*Gucy1b1*^flox/flox^*Ldlr*^-/-^ | Yes (>0.10) | Unpaired t-test |
| S2, D | Body weight, 0 weeks and 10 weeks western diet; *Ldlr*^-/-^ vs. Pf4-*Cre*^+^*Gucy1b1*^flox/flox^*Ldlr*^-/-^ | No: PF4-*Cre*^+^*Gucy1b3*^flox/flox^*Ldlr*^-/-^ 10 weeks (<0.01) | Mann-Whitney test |
| S3, A | Adherent Ly6C^high^ leukocytes; *Ldlr*^-/-^ vs. Pf4-*Cre*^+^*Gucy1b1*^flox/flox^*Ldlr*^-/-^ | Yes (>0.10) | Unpaired t-test |
| S3, B | Adherent LyG^+^ leukocytes; *Ldlr*^-/-^ vs. Pf4-*Cre*^+^*Gucy1b1*^flox/flox^*Ldlr*^-/-^ | Yes (>0.10) | Unpaired t-test |
| S4 | total leukocytes – manual count, FACS; *Ldlr*^-/-^ vs. Pf4-*Cre*^+^*Gucy1b1*^flox/flox^*Ldlr*^-/-^ | Yes (>0.10) | Unpaired t-test |
|  | Ly6G+ leukocytes, FACS; *Ldlr*^-/-^ vs. Pf4-*Cre*^+^*Gucy1b1*^flox/flox^*Ldlr*^-/-^ | Yes (>0.10) |  |
|  | Ly6Chigh monocytes, FACS; *Ldlr*^-/-^ vs. Pf4-*Cre*^+^*Gucy1b1*^flox/flox^*Ldlr*^-/-^ | Yes (>0.10) |  |
|  | Ly6Clow monocytes, FACS; *Ldlr*^-/-^ vs. Pf4-*Cre*^+^*Gucy1b1*^flox/flox^*Ldlr*^-/-^ | Yes (>0.10) |  |
| S5 | *Vcam1* expression, qPCR; WT vs. Pf4-*Cre*^+^*Gucy1b1*^flox/flox^ | Yes (0.07 (WT), >0.10 (KO)) | Paired t-test |
| S6 | Platelet Angpt1 release, ELISA; WT vs. Pf4-*Cre*^+^*Irag1*^flox/flox^ | Yes (>0.10) | Unpaired t-test |
| S8, A | Serum cholesterol, after western diet, *Ldlr*^-/-^; 0 vs. 150 ppm BAY-747 | Yes (>0.10) | Unpaired t-test |
| S8, B | Platelets, after western diet, *Ldlr*^-/-^; 0 vs. 150 ppm BAY-747 | Yes (>0.10) | Unpaired t-test |
| S8, C | Blood leukocytes, after western diet, *Ldlr*^-/-^; 0 vs. 150 ppm BAY-747 | Yes (>0.10) | Unpaired t-test |
| S8, D | Body weight, after western diet, *Ldlr*^-/-^; 0 vs. 150 ppm BAY-747 | Yes (>0.10) | Unpaired t-test |
| S9 | total leukocytes – manual count, FACS; *Ldlr*^-/-^; 0 vs. 150 ppm BAY-747 | Yes (>0.10) | Unpaired t-test |
|  | Ly6G+ leukocytes, FACS; *Ldlr*^-/-^; 0 vs. 150 ppm BAY-747 | Yes (>0.10) | Unpaired t-test |
|  | Ly6Chigh monocytes, FACS; *Ldlr*^-/-^; 0 vs. 150 ppm BAY-747 | Yes (>0.10) | Unpaired t-test |
|  | Ly6Clow monocytes, FACS; *Ldlr*^-/-^; 0 vs. 150 ppm BAY-747 | Yes (>0.10) | Unpaired t-test |

**Suppl. Table S2**: Characteristics of individuals enrolled in the analysis of angiopoietin-1 release in activated platelets. Unless otherwise indicated, data are mean and s.e.m. * Unpaired t-test, ^†^ Chi-square test. *PRP*, platelet rich plasma.

| **Characteristic** | **Genotype** | | **p-value** |
| --- | --- | --- | --- |
|  | AA/AG  N=5 | GG  N=5 |  |
| Age, years | 37.4 ± 5.3 | 30.8 ± 1.8 | 0.27* |
| Female gender, n (%) | 4 (80) | 4 (80) | >0.99^†^ |
| Presence of CAD, n (%) | 0 (0) | 0 (0) | >0.99^†^ |
| Antiplatelet medication, n (%) | 0 (0) | 0 (0) | >0.99^†^ |
| PRP platelet count, 10^3^ platelets | 365.8 ± 52.2 | 261.0 ± 18.2 | 0.09* |

**Suppl. Table S3**: Enriched pathways in co-expression module 11.

| **ID** | **Name** | **Bonferoni p** |
| --- | --- | --- |
| GO:0009611 | response to wounding | 4.71·10^-63^ |
| GO:0042060 | wound healing | 1.84·10^-60^ |
| GO:0050878 | regulation of body fluid levels | 9.80·10^-49^ |
| GO:0007596 | blood coagulation | 1.99·10^-48^ |
| GO:0050817 | coagulation | 3.23·10^-48^ |
| GO:0007599 | hemostasis | 4.44·10^-48^ |
| GO:0046903 | secretion | 2.28·10^-46^ |
| GO:0007167 | enzyme linked receptor protein sign. pathway | 6.11·10^-44^ |
| GO:0040012 | regulation of locomotion | 6.64·10^-44^ |
| GO:2000145 | regulation of cell motility | 1.31·10^-43^ |
